# Amino acids stimulate the endosome-to-Golgi trafficking through Ragulator and small GTPase Arl5

**DOI:** 10.1101/312546

**Authors:** Meng Shi, Bing Chen, Boon Kim Boh, Yan Zhou, Divyanshu Mahajan, Hieng Chiong Tie, Lei Lu

## Abstract

The endosome-to-Golgi or endocytic retrograde trafficking pathway is an important post-Golgi recycling route. We made a novel discovery that the retrograde trafficking of cargos is inhibited and stimulated by the absence and presence, respectively, of amino acids (AAs), especially glutamine. By testing components of the AA-stimulated mTORC1 signaling pathway, we demonstrated that SLC38A9, v-ATPase and Ragulator, but not Rag GTPases and mTORC1, are essential for the AA-stimulated trafficking. Arl5, an ARF-like family small GTPase, interacts with Ragulator in an AA-regulated manner and both Arl5 and its effector, the Golgi-associated retrograde protein complex (GARP), are required for the AA-stimulated trafficking. We have therefore identified a mechanistic connection between the nutrient signaling and the retrograde trafficking pathway, whereby SLC38A9 and v-ATPase sense AA-sufficiency and Ragulator functions as a guanine nucleotide exchange factor to activate Arl5, which, together with GARP, a tethering factor, probably facilitates the endosome-to-Golgi trafficking.

## Introduction

In eukaryotic cells, proteins and lipids (cargos) are dynamically exchanged among cellular organelles through trafficking routes or pathways. In the endocytic pathway, cargos on the plasma membrane (PM) are internalized to the early endosome (EE), where cargos can have distinct subsequent endocytic fates. For example, the progression of the EE to the late endosome (LE) can lead to the eventual degradation of cargos in the lysosome. Alternatively, the retrograde or the endosome-to-Golgi trafficking pathway diverts internalized cargos to the *trans*-Golgi network (TGN), therefore salvaging cargos from degradation; from the TGN, cargos return to either the PM or the endosome to complete their itinerary cycles(Bonifacino and Rojas, 2006; Burd, 2011; Chia et al., 2013; Johannes and Popoff, 2008; Lu and Hong, 2014; Maxfield and McGraw, 2004). A growing list of cargos, including most TGN resident transmembrane proteins (TGN membrane proteins), has been documented to take this trafficking route. Pathogens, such as viruses and plant or bacterial toxins, can also hijack this pathway to invade cells while avoiding lysosomal degradation. As a major cellular recycling pathway, the endosome-to-Golgi trafficking has been recognized for the post-Golgi secretion, biogenesis of the lysosome, maintenance of sphingolipid homeostasis, regulation of Wnt signaling and pathogenesis of neurodegenerative diseases(Frohlich et al., 2015; Hirata et al., 2015; Perez-Victoria et al., 2008).

Recent progress in this field offers us a rough picture on how the endosome-to-Golgi trafficking works at molecular and cellular level(Bonifacino and Rojas, 2006; Chia et al., 2013; Johannes and Popoff, 2008; Lu and Hong, 2014; Pfeffer, 2011). First, cargos are selected and sorted into a membrane carrier at the endosomal membrane in conjunction with clathrin coats, coat adaptors, retromer, GARP (Golgi-associated retrograde protein complex) and actin cytoskeleton. Next, the budded membrane carrier is targeted to the TGN by microtubule motor. After that, with the aid of small GTPases such as Arl1 and Rab6, the carrier attaches to the TGN membrane by tethering factors such as Golgins and GARP. Finally, the formation of SNARE complex drives the fusion between the carrier and the TGN membrane, accomplishing the delivery of cargos to the Golgi.

Nutrients, including amino acids (AAs), glucose and other carbon sources, are the most fundamental resources for the growth and proliferation of cells. Nutrient sufficiency stimulates anabolic metabolism, such as the biosynthesis of macromolecules and the biogenesis of organelles; on the other hand, nutrient starvation triggers catabolic pathways, such as autophagy, to break down macromolecules in order to recycle much needed materials for cell survival. Cells must have evolved sophisticated signaling networks to coordinate their sub-cellular activities according to the environment nutrient. In yeast, it is known that the amount of AA-permeases on the PM is regulated by AAs. Yeast Gap1p (general AA permease 1) is one of the most studied cargos for the nutrient-regulated endocytosis. In response to AA-starvation, Gap1p accumulates on the PM to scavenge extracellular nitrogen sources; whereas in the presence of AAs, especially Gln, TORC1 is activated to initiate a signaling cascade to internalize and degrade Gap1p in the vacuole(Chen and Kaiser, 2002; Lauwers et al., 2009; Merhi and Andre, 2012; Roberg et al., 1997). The best known cargo undergoing AA-regulated intracellular membrane trafficking is probably Atg9, a conserved transmembrane protein essential for autophagy. AA-starvation induces the translocation of Atg9 from the peripheral pool to the phagophore assembly site in yeast(Mari et al., 2010) or from the Golgi to the endosome via Ulk1-dependent signaling pathway in mammalian cells(Young et al., 2006). Besides Atg9, it is currently unknown in mammalian cells if and how nutrient regulates intracellular membrane trafficking, especially the endosome-to-Golgi pathway.

In contrast, a great deal of molecular details have been known on how AAs regulate cellular metabolism through transcription and translation. The cell’s decision on anabolic or catabolic metabolism is mainly made through the mechanistic target of rapamycin complex 1 (mTORC1) signaling pathway, which senses the presence of nutrients and growth factors in combination with the cellular energy and stress status(Efeyan et al., 2012; Jewell and Guan, 2013; Shimobayashi and Hall, 2014). When extracellular AAs are abundant, there is a rapid accumulation of AAs within the lumen of the lysosome. Luminal AAs trigger SLC38A9(Jung et al., 2015; Rebsamen et al., 2015; Wang et al., 2015), a SLC-family AA transceptor, and vacuolar adenosine triphosphatase (v-ATPase)(Zoncu et al., 2011), a proton pump responsible for the acidification of the lysosome. Next, activated SLC38A9 and v-ATPase signal to Ragulator by rearranging their interaction with the latter. Ragulator is a pentameric complex comprising Lamtor1-5, which are also referred to as p18, p14, MP1, C7orf59 and HBXIP, respectively(Bar-Peled et al., 2012; Sancak et al., 2010). Following the activation, Ragulator functions as the guanine nucleotide exchange factor (GEF) of heterodimeric Rag GTPases(Bar-Peled et al., 2012). Finally, GTP-loaded Rag heterodimer recruits mTORC1 to the lysosomal surface(Sancak et al., 2010), where the full kinase activity of mTORC1 is turned on by growth factor-activated small GTPase, Rheb(Long et al., 2005). Active mTORC1 initiates anabolic processes through translation and transcription by phosphorylating a cascade of its substrates.

Here we investigated if nutrients can regulate the endosome-to-Golgi trafficking and discovered that the trafficking is impeded by AA-starvation and promoted by the presence of AAs, especially Gln. We subsequently identified a mechanistic connection between the AA-sensing module of the mTORC1 signaling and retrograde trafficking. We propose that, in the AA-stimulated retrograde trafficking, SLC38A9 and v-ATPase sense AAs and activate Ragulator, which functions as a GEF for Arl5, an ARF-like (Arl) family small GTPase; active Arl5 probably recruits GARP to facilitate the delivery of cargos to the TGN.

## Results

### Starvation reversibly induces the translocation of TGN membrane proteins to the endosomal pool

To investigate if nutrient plays a role in the endocytic membrane trafficking, we compared the sub-cellular distribution of TGN membrane proteins in normal and starvation medium. Most TGN membrane proteins, such as furin, cation-independent mannose 6-phosphate receptor (CI-M6PR), cation-dependent mannose 6-phosphate receptor (CD-M6PR) and sortilin, cycle between the PM and TGN through endosomes(Lu and Hong, 2014). Their relative distribution between the Golgi and endosomal pool shifts in response to a change in the endocytic trafficking. In the complete medium (DMEM supplemented with 10% fetal bovine serum), endogenous furin mainly colocalized with Golgin-245 at the TGN in HeLa cells, as expected (Fig. 1a,b). When serum or growth factor was withdrawn by incubation in DMEM for 1 h, no significant change of furin was observed (Fig. 1a,b). In contrast, when cells were starved of both AAs and growth factors by incubating in Hank’s balanced salt solution (HBSS) for 1 h, furin lost its Golgi-pool and appeared diffused throughout the cytosol (Fig. 1a,b). The starvation-induced translocation was reversible. When HBSS-treated cells were subsequently supplied with nutrient (DMEM), furin rapidly re-appeared at the Golgi and recovered to pre-starvation state after 1 h (Fig. 1c). The finding was also observed using exogenously expressed full length furin-GFP (Fig. 1d and Supplementary Fig. 1a). The rapid and reversible distribution demonstrated that furin was probably not degraded and, instead, it was likely arrested at the peripheral pool.

**Figure 1.**
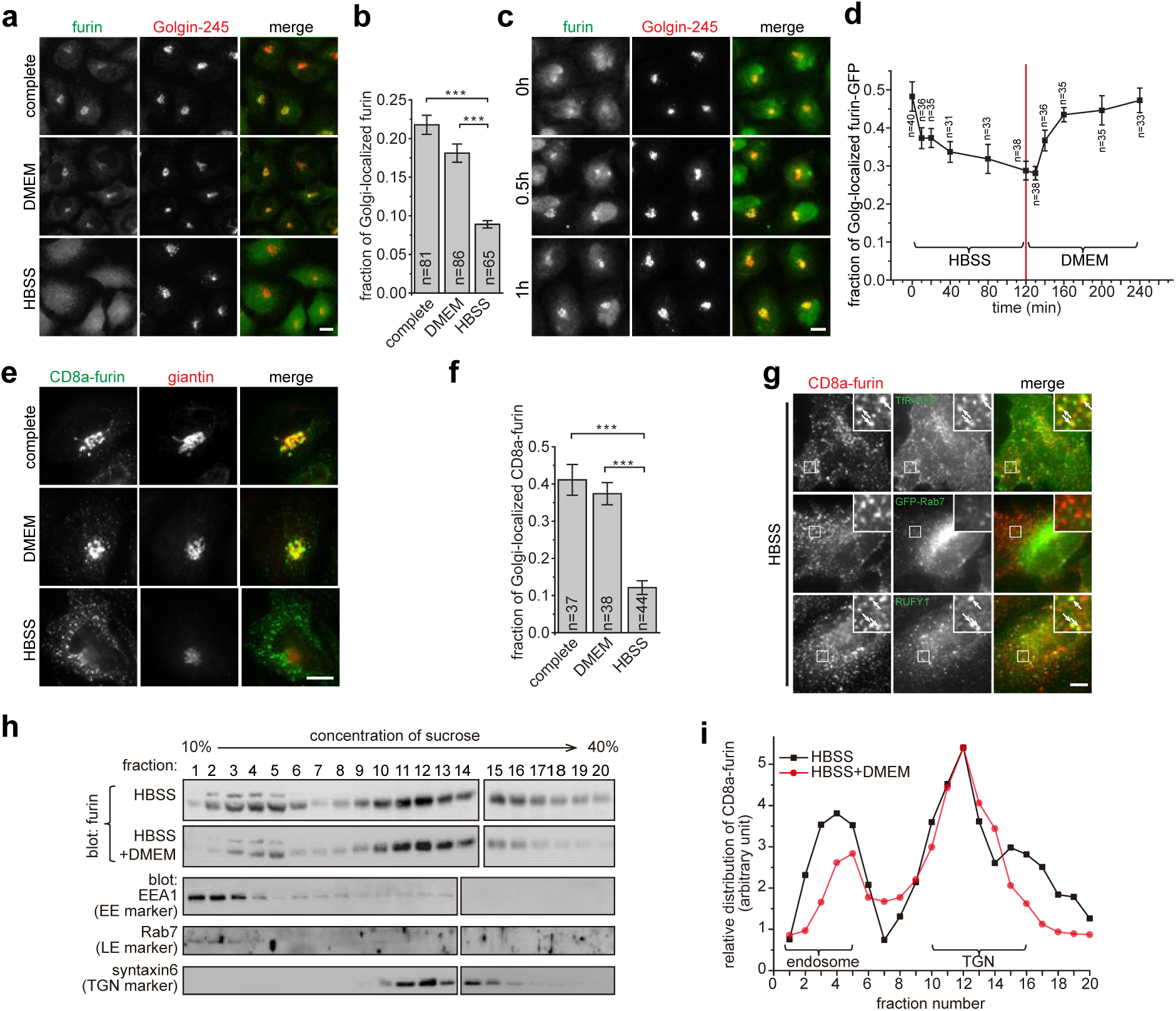
Nutrient starvation reversibly induces the translocation of TGN membrane proteins to the endosomal pool. All cells are HeLa cells. (**a,b**) Furin loses its Golgi localization during starvation. Cells treated with indicated medium for 1 h and endogenous furin and Golgin-245 were stained. The fraction of Golgi-localized furin is quantified in **b**. (**c**) The recovery of Golgi localization of furin after supplying nutrient. After starvation in HBSS for 2 h, cells were treated with DMEM for indicated time. Endogenous furin and Golgin-245 were stained. (**d**) Kinetics of Golgi-localized furin-GFP during HBSS and subsequent DMEM treatment. Cells expressing furin-GFP were first starved in HBSS for 2 h and subsequently stimulated by DMEM for 2 h. At indicated time, cells were stained for endogenous giantin and the fraction of Golgi-localized furin-GFP is quantified. (**e,f**) Nutrient starvation significantly reduces the Golgi localization of CD8a-furin. Cells transiently expressing CD8a-furin were treated by indicated medium for 2 h and CD8a-furin and endogenous giantin were stained in **e**. The fraction of Golgi-localized CD8a-furin is quantified in **f**. (**g**) The translocation of CD8a-furin to the endosome during nutrient starvation. Cells transiently expressing CD8a-furin singly or together with indicated marker constructs were treated with HBSS for 2 h and stained to reveal indicated proteins. Boxed regions are enlarged at the upper right corner. Arrows indicate colocalization. (**h,i**) The endosomal fraction of CD8a-furin increases during nutrient starvation. Similar results have been observed in four independent experiments. Cells stably expressing CD8a-furin were treated with HBSS for 2 h or with additional 20 min treatment of DMEM. Lysates were subjected to sucrose gradient centrifugation to separate organelles. 20 fractions were collected and immuno-blotted to locate the distribution of CD8a-furin and markers. The distribution of CD8a-furin is quantified in **i**. Two curves are normalized by their peaks at fraction 12. Complete, complete medium; n, the number of cells analyzed; error bar, s.e.m.; scale bar, 10 μm. *P*-values were from *t*-test. ***, *P* ≤ 0.0005.

To further confirm our observation and resolve the peripheral location of furin, we imaged exogenously expressed CD8a-furin chimera, which consists of CD8a luminal and transmembrane domain and furin cytosolic domain(Mahajan et al., 2013). We similarly observed the reversible change of CD8a-furin localization under starvation and nutrient stimulation condition (Fig. 1e,f). Under starvation, the peripheral pool of CD8a-furin colocalized with the EE marker, RUFY1(Cormont et al., 2001), and the recycling endosome (RE) marker, transferrin receptor-GFP (TfR-GFP)(Gruenberg and Maxfield, 1995), but not the LE marker, GFP-Rab7(Chavrier et al., 1990) (Fig. 1g), suggesting that it mainly localized to the EE and RE during starvation. The colocalization study using full length furin confirmed these results, although a significant localization to the LE was also observed (Supplementary Fig. 1b), probably due to the contribution of native transmembrane domain(Chia et al., 2011). The preferential endosomal distribution of CD8a-furin under starvation was also biochemically demonstrated using sucrose gradient ultracentrifugation (Fig. 1h). Endosomal and TGN fractions were identified by EEA1 and syntaxin6, respectively(Bock et al., 1997; Mu et al., 1995) (Fig. 1h). From quantitative Western blot, endosomal fractions of starved cells contained more CD8a-furin than those of AA-stimulated cells (Fig. 1i).

Starvation-induced change of localization was also similarly observed for other TGN membrane proteins such as endogenous CI-M6PR and overexpressed CD8a-CI-M6PR (Supplementary Fig. 1c-f), though the effect varied with respect to proteins. Our subsequent studies mainly focused on CD8a-furin since its subcellular localization seems to be most sensitive to nutrient among TGN membrane reporters that we tested.

### AAs but not glucose and growth factors stimulate the endosome-to-Golgi trafficking

The reduction of the Golgi pool and the concomitant increase in the endosomal pool under HBSS treatment suggest that DMEM and complete medium might stimulate the endosome-to-Golgi trafficking. The complete medium used in our cell culture comprised serum, which is the source of growth factors, and DMEM, which consists of DMEM-base (inorganic salts and vitamins), AAs (15 AAs including Gln) and glucose. To reveal the component(s) behind the stimulation, the PM-to-Golgi trafficking assay was conducted in the testing medium comprising DMEM-base supplemented with combinations of dialyzed-serum, AAs and glucose. To that end, CD8a-furin expressing cells were first starved in DMEM-base. The surface-exposed CD8a-furin was subsequently labeled and the antibody-CD8a-furin complex was monitored along the endocytic trafficking en route to the Golgi. We found that AAs, but not glucose or serum, were sufficient to stimulate the endocytic trafficking of CD8a-furin to the Golgi (Fig. 2a,b). Supplementation of AAs, dialyzed-serum and glucose in DMEM-base, or the usage of the complete medium, produced no more stimulatory effect than AAs alone (Fig. 2a,b). In fact, a weaker stimulation by the complete medium than AAs was often observed, suggesting an unknown adverse effect of glucose and growth factors on the endocytic trafficking. Similar AA-stimulation effect was also observed in BSC-1 cells or by using other reporters, such as CD8a-fused CI-M6PR, CD-M6PR and sortilin, in HeLa cells (Supplementary Fig. 2a,b), demonstrating that the effect of AAs is probably ubiquitous.

**Figure 2.**
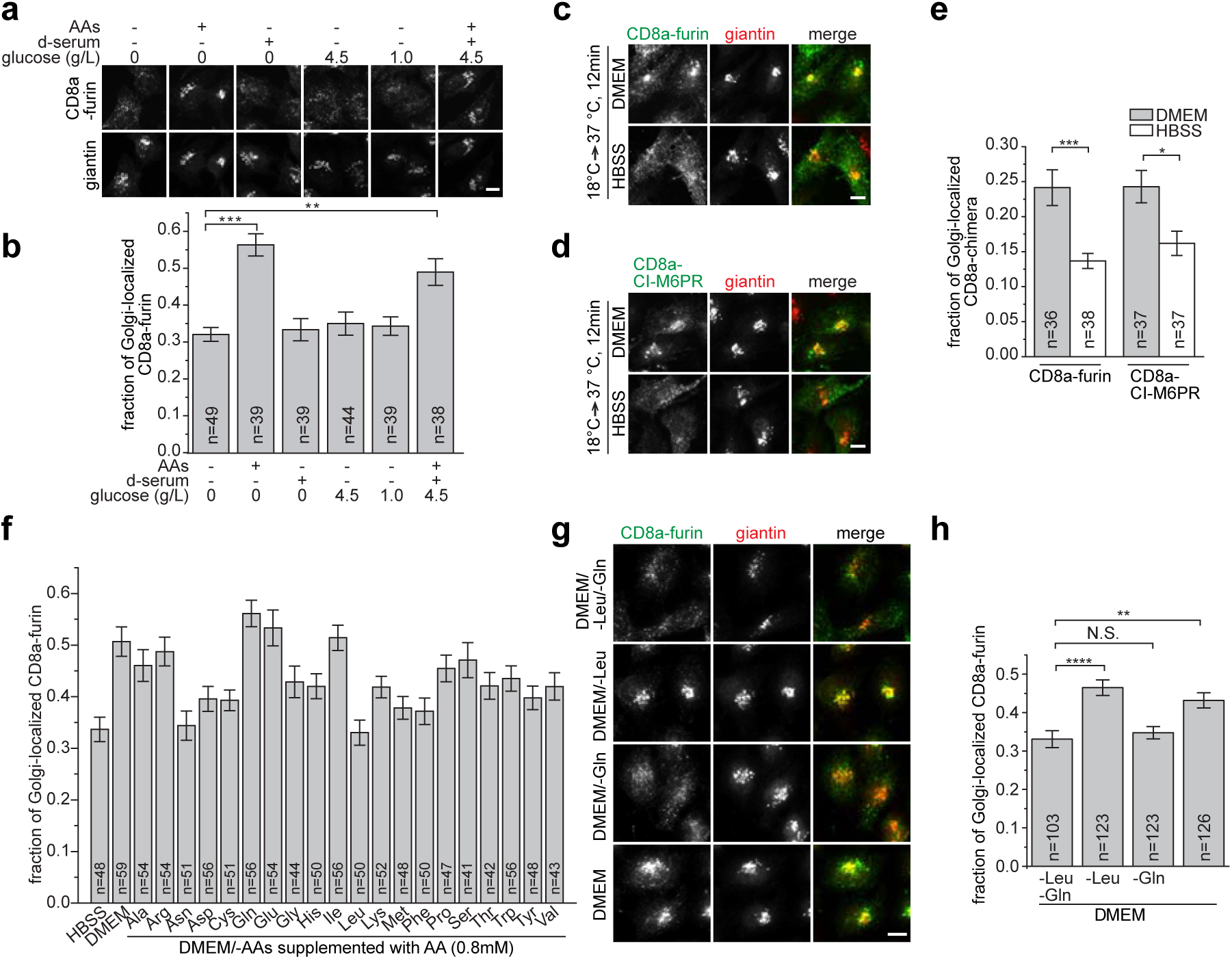
AAs stimulate the endosome-to-Golgi trafficking. All cells are HeLa cells. (**a**,**b**) AAs but not growth factors and glucose stimulate the retrograde trafficking to the Golgi. Cells stably expressing CD8a-furin were treated with DMEM-base for 2 h. The surface-exposed CD8a-furin was labeled by anti-CD8a antibody and chased in respective medium for 20 min. After staining CD8a and endogenous giantin, the fraction of CD8a-furin at the Golgi is quantified. d-serum, 10% dialyzed serum. (**c-e**) AAs stimulate the endosome-to-Golgi trafficking. Cells transiently expressing CD8a-furin or CD8a-CI-M6PR were surfaced-labeled by anti-CD8a antibody and synchronized at endosomes at 18 °C in HBSS for 2 h before being chased at 37 °C in HBSS or DMEM for 12 min. CD8a and endogenous giantin were stained and the fraction of Golgi-localized CD8a chimeras is quantified. (**f-h**) Gln has one of the most acute stimulating effects on endocytic trafficking to the Golgi. Similar to **a** and **b** except that the nutrient starvation was conducted in HBSS before surface-labeling. The labeled CD8a-furin was then chased in HBSS, DMEM, DMEM/-AAs supplemented with indicated AA at 0.8 mM in **f** or DMEM selectively leaving out indicated AA(s) in **g**. Data shown in **f** is representative of three independent experiments. The fraction of Golgi-localized CD8a-furin is quantified in **h**. n, the number of cells analyzed; error bar, s.e.m.; scale bar, 10 μm. *P*-values were from *t*-test. N.S., not significant (*P* > 0.05); *, *P* ≤ 0.05; **, *P* ≤ 0.005; ***, *P* ≤ 0.0005; ****, *P* ≤ 0.00005.

The endocytic trafficking of CD8a-furin to the Golgi comprises two consecutive steps, the clathrin-dependent endocytosis from the PM to the endosome and the subsequent endosome-to-Golgi or the retrograde trafficking. Our quantitative analysis indicated that the internalized CD8a-furin within 6 min of chase did not display an obvious difference between AA-sufficiency and-starvation (Supplementary Fig. 2d), therefore suggesting that the endocytosis could not be the target of AA-stimulation. Hence, we directly tested the retrograde trafficking. Antibody-labeled CD8a-furin or -CI-M6PR was first allowed to accumulate at the EE and RE using 18 °C synchronization protocol(Mallard et al., 1998; Tai et al., 2004). The Golgi localization was subsequently quantified after a chase at 37 °C for 12 min in the medium with or without AAs. We found that AAs significantly increased the Golgi-localized CD8a-furin and-CI-M6PR (Fig. 2c-e). In summary, our data demonstrate that AAs stimulate the endosome-to-Golgi trafficking of furin and other TGN membrane proteins. Upon AA-starvation, the endosome-to-Golgi trafficking is compromised; as the Golgi-to-PM anterograde trafficking proceeds, an elevated endosomal pool is resulted for the TGN membrane protein at the expense of its Golgi one.

### Glutamine has the most acute effect on stimulating the endosome-to-Golgi trafficking

To determine which AA(s) is(are) responsible for the AA-stimulated endosome-to-Golgi trafficking, we tested each of 20 AAs at the same concentration (0.8 mM) for their effect on the endocytic trafficking. While a majority of AAs stimulated the trafficking to various degrees, Gln, a non-essential AA, consistently stood out as the most potent stimulator (Fig. 2f). The stimulating effect of Gln was concentration dependent and peaked at ~ 0.6 mM (Supplementary Fig. 2e). Other stimulatory AAs, such as Ala, followed similar trend. To our surprise, Leu, which is an essential AA and the most acute stimulator for mTORC1 signaling(Lynch, 2001), as well as Asn, displayed minimal stimulation (Fig. 2f). The lack of stimulation by Leu was probably not due to suboptimal concentration used in our assay since similar result was observed for a wide range of concentrations from 0.01 to 5.12 mM (Supplementary Fig. 2e). We found that the effect of combining different AAs appeared complex and was not simply additive (Supplementary Fig. 2f), the mechanism behind which awaits future exploration. The observation is nonetheless consistent with the finding that DMEM, which contains 15 AAs, repeatedly showed less activity than Gln alone (Fig. 2f). It is possible that AAs can positively or negatively modulate each other’s activities, considering that certain AAs can act as exchangers for others(Krall et al., 2016).

To further test if Gln is essential for the retrograde trafficking in parallel comparison with Leu, we investigated the subcellular distribution of CD8a-furin in DMEM selectively leaving out Gln (DMEM/-Gln), Leu (DMEM/-Leu) or both (DMEM/-Leu/-Gln) (Fig. 2g,h). In media without Gln (DMEM/-Leu/-Gln and DMEM/-Gln), CD8a-furin mainly localized to peripheral puncta at the expense of its Golgi localization, indicating an essential role of Gln in the retrograde trafficking; in media containing Gln (DMEM/-Leu and DMEM), CD8a-furin prominently accumulated in the Golgi. In contrast, the depletion of Leu (DMEM/-Leu) did not affect the Golgi localization of CD8a-furin comparing with DMEM. We ruled out the possibility that our HeLa cells became insensitive to Leu since, consistent with our current knowledge(Chiu et al., 2012; Duran et al., 2012; Jewell et al., 2015; Nicklin et al., 2009), Gln was observed to be essential and synergize with Leu for the activation of mTORC1 signaling (Supplementary Fig. 2g). Collectively, we conclude that Gln, but not Leu, is a necessary and sufficient stimulator for the endosome-to-Golgi trafficking.

### The AA-stimulated retrograde trafficking depends on v-ATPase, SLC38A9 and Ragulator but not Rag GTPases and mTORC1

It is known that nutrient signaling culminates in the activation of mTORC1 through SLC38A9, v-ATPase, Ragulator and heterodimeric Rag GTPases(Bar-Peled et al., 2012; Jung et al., 2015; Rebsamen et al., 2015; Sancak et al., 2010; Wang et al., 2015; Zoncu et al., 2011). To test if AA-stimulated retrograde trafficking utilizes a similar pathway, we selectively compromised each component through small molecule inhibitors or RNAi-mediated knockdowns and subsequently investigated the ensuing effect on the retrograde trafficking. To compare and contrast the stimulatory effect, the fraction of Golgi-localized CD8a-furin under AA-sufficiency was normalized by that under AA-starvation to yield a quantity referred to as the AA-stimulated Golgi trafficking. In the presence of concanamycin A (conA), an inhibitor of v-ATPase(Bowman et al., 2004), CD8a-furin was arrested at peripheral endosomes even under AA-sufficiency (Supplementary Fig. 3a). The AA-stimulated Golgi trafficking decreased significantly under conA, in comparison with the control (Fig. 3a). When SLC38A9, a high affinity transporter for Gln(Rebsamen et al., 2015), was depleted by its shRNAs (Fig. 3b), the AA-stimulated mTORC1 activity was significantly reduced as previously expected (Jung et al., 2015; Rebsamen et al., 2015; Wang et al., 2015) (Fig. 3c) and so did AA-stimulated Golgi trafficking (Fig. 3d). Similarly, when Lamtor1 and Lamtor3, two subunits of Ragulator, were individually depleted by corresponding shRNAs (Fig. 3e; Supplementary Fig. 3b), the AA-stimulated Golgi trafficking decreased significantly in comparison with the control (Fig. 3f), suggesting that intact Ragulator is required for the AA-stimulated retrograde trafficking.

**Figure 3.**
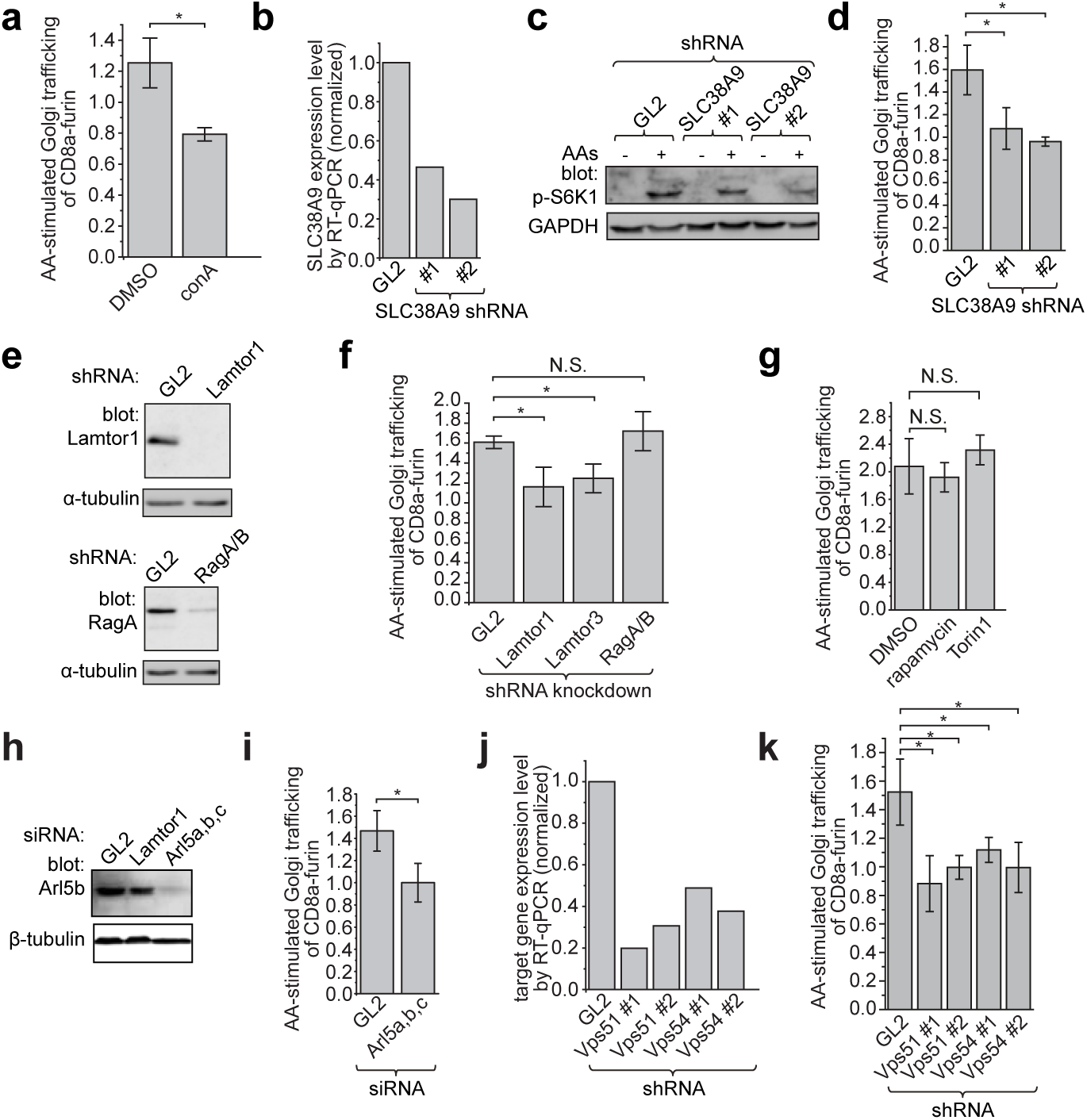
v-type ATPase, SLC38A9, Ragulator, Arl5 and GARP, but not mTORC1 and Rag GTPases, are essential for the AA-stimulated endosome-to-Golgi trafficking of CD8a-furin. (**a**) HeLa cells stably expressing CD8a-furin were starved in HBSS for 2 h followed by surface-labeling and subsequent incubation with either HBSS or DMEM for 20 min. 1% DMSO or 2.5 μM conA was present throughout the incubation. Cells were stained and the AA-stimulated Golgi trafficking is quantified by imaging. (**b**) Endogenous SLC38A9 was knocked down by lentivirus-transduced shRNAs as assessed by RT-qPCR. (**c**) The knockdown of endogenous SLC38A9 attenuated the AA-stimulated mTORC1 activity. SLC38A9 knockdown cells were incubated with DMEM/-AAs for 50 min followed by incubation with DMEM for 20 min. Cell lysates were immuno-blotted for p-S6K1 and GAPDH. (**d**) SLC38A9 is required for the AA-simulated Golgi trafficking. Cells knocked down by indicated shRNAs were transfected to express CD8a-furin and subjected to treatment and analysis similar to **a**. (**e**) Immuno-blots showing that endogenous Lamtor1 and RagA/B were knocked down by respective lentivirus-transduced shRNAs. (**f**) Lamtor1 and 3 but not RagA/B are required for the AA-stimulated Golgi trafficking. The experiment was conducted similarly to **d**. (**g**) mTORC1 is not required for the AA-stimulated retrograde trafficking. The experiment was conducted similarly to **a** except that 1% DMSO, 100 nM rapamycin or 250 nM Torin1 was present throughout the treatment. (**h**) The immuno-blot showing that endogenous Arl5b was knocked down by a mixture of siRNAs targeting Arl5a, b and c. (**i**) Arl5 is required for the AA-stimulated Golgi trafficking of CD8a-furin. Experiment was conducted similarly to **d**. (**j**) The knockdown of endogenous Vps51 and Vps54 by respective lentivirus-transduced shRNAs as assessed by RT-qPCR. (**k**) GARP is required for the AA-stimulated Golgi trafficking. Experiment was conducted similarly to **d**. In **a**, **d**, **f**, **g**, **I** and **k**, the displayed value is the mean of n=3 independent experiments with each experiment analyzing ≥ 90, 21, 61, 50, 51 and 31 cells respectively; error bar, s.d.. *P*-values were from *t*-test. N.S., not significant (*P* > 0.05); *, *P* ≤ 0.05. GL2 is a non-targeting control siRNA or shRNA.

Our findings on the AA-stimulated trafficking so far seem consistent with the AA-stimulated mTORC1 signaling pathway, in which SLC38A9, v-ATPase, Ragulator and Rag GTPases act sequentially in recruiting mTORC1 to the lysosome(Efeyan et al., 2012; Jewell and Guan, 2013; Shimobayashi and Hall, 2014). However, simultaneous depletion of both Rag A and B by shRNAs (Fig. 3e,f), a condition in which AA-stimulated mTORC1 activation was inhibited (Supplementary Fig. 3c), did not significantly affect the AA-stimulated Golgi trafficking (Fig. 3f). Lastly, when mTORC1 activity was strongly inhibited by rapamycin or Torin1 (Supplementary Fig. 3d), the AA-stimulated Golgi trafficking did not significantly change either (Fig. 3g). Altogether, our results demonstrated that SLC38A9, v-ATPase and Ragulator, but not Rag GTPases and mTORC1, are probably involved in the AA-stimulated endosome-to-Golgi trafficking.

### Arl5b interacts with Ragulator through Lamtor1

The endosome-to-Golgi trafficking requires Rab and Arl-family small GTPases(Bonifacino and Rojas, 2006; Johannes and Popoff, 2008; Lu and Hong, 2014; Pfeffer, 2011) and we previously discovered Arl1 as a key regulator for this pathway(Lu et al., 2001; Lu et al., 2004). In our quest for additional Arl regulators for this pathway, we focused on Arl5b, which resides on the Golgi and regulates the endosome-to-Golgi and the reverse trafficking, probably by interacting with its effector GARP and AP4, respectively(Houghton et al., 2012; Rosa-Ferreira et al., 2015; Toh et al., 2017). To gain insight into the upstream regulators and downstream effectors of Arl5b, we performed a yeast two-hybrid screen of human kidney cDNA library using GTP-bound mutant form of Arl5b as the bait. Interestingly, one of the strongest hits identified was full length Lamtor1, a subunit of Ragulator.

The interaction between Arl5b and Lamtor1 was first biochemically confirmed in pull-down and immunoprecipitation assays. Bead-immobilized GST-Arl5b and a control Arl, GST-Arl1, were in vitro loaded with guanosine 5′-[β,γ-imido]triphosphate (GMPPNP; a non-hydrolyzable GTP analog) and GDP. The resulting beads were then incubated with 293T cell lysate expressing Lamtor1-GFP. Both GMPPNP and GDP-loaded Arl5b pulled down Lamtor1-GFP (Fig. 4a). In contrast, neither GMPPNP nor GDP-loaded Arl1 retained Lamtor1-GFP (Fig. 4a), demonstrating the specificity of the interaction between Arl5b and Lamtor1. Similar to Arl1 and other Ras-family small GTPases(Lu et al., 2001), guanine nucleotide binding mutations, Q70L (GTP-bound mutant form; hereafter QL) and T30N (GDP-bound mutant form; hereafter TN), were introduced to Arl5b to make it constitutively active and inactive, respectively(Houghton et al., 2012). In 293T cell lysates co-expressing C-terminally GFP-tagged Arl5b-wild type (hereafter wt), QL, TN or Lamtor2 together with Lamtor1-Myc or SNX3-Myc, we found that Arl5b-QL and-TN, but not Arl5b-wt, interacted with Lamtor1 in both forward and reverse co-immunoprecipitations (co-IPs) (Fig. 4b,c). As previously reported(Bar-Peled et al., 2012; Sancak et al., 2010), exogenously expressed Lamtor2-GFP interacted with Lamtor1-Myc (Fig. 4b,c).

**Figure 4.**
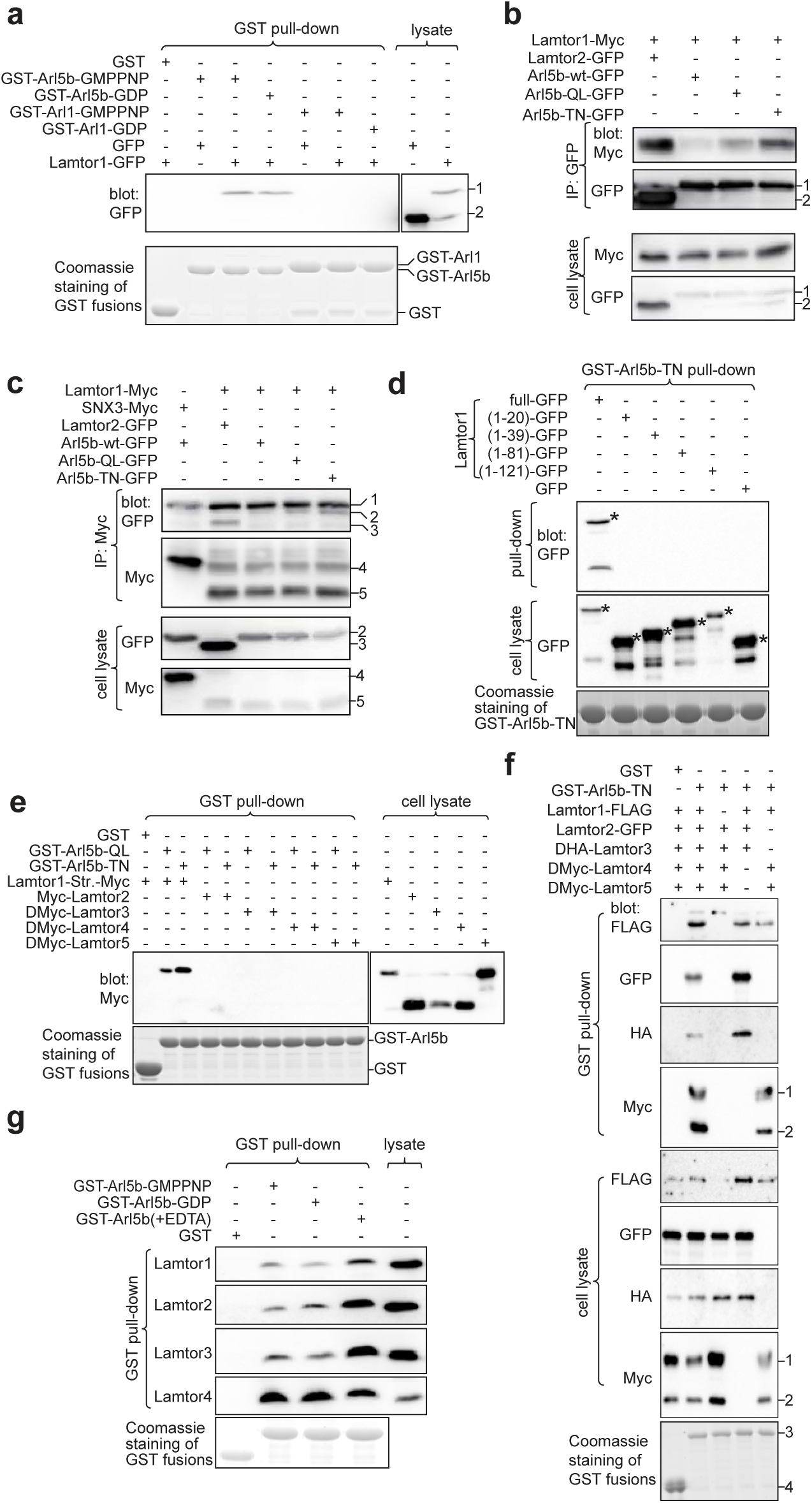
Arl5b interacts with Ragulator through Lamtor1. HEK293T cells were used. (**a**) Arl5b, but not Arl1, specifically pulled down Lamtor1-GFP. Bead-immobilized GST-Arl5b or Arl1 was *in vitro* loaded with GDP or GMPPNP and subsequently incubated with cell lysates expressing GFP or Lamtor1-GFP. Pull-downs were analyzed by immuno-blotting GFP-fusions. The loading of GST-fusions were shown by Coomassie blue staining. “1” and “2” indicate Lamtor1-GFP and GFP band, respectively. (**b,c**) Arl5b-QL and-TN interact with Lamtor1-Myc in forward and reverse co-IPs. Cells co-expressing indicated tagged-proteins were incubated with indicated antibodies and IPs were immuno-blotted against indicated tags. Lamtor2-GFP and SNX3-Myc served as a positive and negative control, respectively. In **b**, “1” and “2” indicate Arl5b-(wt, QL or TN)-GFP and Lamtor2-GFP band, respectively. In **c**, “1-5” indicate IgG heavy chain, Arl5b-(wt, QL or TN)-GFP, Lamtor2-GFP, SNX3-Myc and Lamtor1-Myc band, respectively. **(d)** Full length Lamtor1 is required for Arl5b-Lamtor1 interaction. Bead-immobilized GST-Arl5b-TN was incubated with cell lysates expressing indicated fragments of Lamtor1 and pull-downs were analyzed by immuno-blotting GFP-fusions. “*” denotes the specific band. (**e**) Arl5-QL and-TN interact with Lamtor1 but not Lamtor2-5. Bead-immobilized GST-fusion was incubated with cell lysates expressing indicated Myc-tagged Lamtors and pull-downs were analyzed by immuno-blotting Myc-tagged proteins. Lamtor1-Str.-Myc, Lamtor1-G2A-Strep-Myc. (**f**) Arl5b-TN interacts with Ragulator through Lamtor1. Bead-immobilized GST-Arl5b-TN was incubated with cell lysates expressing indicated Lamtors and pull-downs were analyzed by immuno-blotting indicated tags. “1-4” indicate DMyc-Lamtor5, DMyc-Lamtor4, GST-Arl5b-TN and GST band, respectively. (**g**) Arl5b-GMPPNP,-GDP and guanine nucleotide empty form interact with Ragulator. Bead-immobilized GST-Arl5b were first loaded with GMPPNP or GDP or stripped off its bound guanine nucleotide by EDTA treatment. Beads were subsequently incubated with cell lysate and pull-downs were analyzed by immuno-blotting endogenous Lamtor1-4.

To investigate Arl5b-binding region of Lamtor1, we generated GFP-tagged serial truncates of Lamtor1. When 293T cell lysates expressing individual truncates were incubated with bead-immobilized GST-Arl5b-TN, it was found that Arl5b-Lamtor1 interaction was abolished by the removal of either N- or C-terminus of Lamtor1, indicating that both ends of Lamtor1 are required for interaction (Fig. 4d). Previous studies have established that Lamtor1 anchors Ragulator to the lysosomal membrane by its N-terminal dual-lipid modification and it functions as a scaffold to independently bind to two heterodimeric sub-complexes—Lamtor2-Lamtor3 and Lamtor4-Lamtor5(Bar-Peled et al., 2012; Nada et al., 2009; Sancak et al., 2010). We characterized the interaction between Arl5b and individual subunits or sub-complexes. Except for Lamtor1, immobilized GST-Arl5b-QL or -TN did not pull down individually expressed Lamtor2-4 (Fig. 4e). When incubated with cell lysates expressing combinations of exogenously expressed Ragulator subunits, immobilized GST-Arl5b pulled down Lamtor2-Lamtor3 and Lamtor4-Lamtor5 sub-complexes only in the presence of co-expressed Lamtor1 (Fig. 4f). In addition to exogenously expressed Lamtors, immobilized GST-Arl5b also pulled down endogenous Lamtors (Fig. 4g). In summary, we conclude that Arl5b interacts with Ragulator through Lamtor1. Although both GTP and GDP-mutant forms interacted with Lamtor1, GDP-mutant form of Arl5b appeared to interact more strongly (Fig. 4b,c,e), the significance of which is discussed later.

In human and mouse genome, there are three paralogs of Arl5, Arl5a, b and c, with AA sequence identity ≥ 64%. In contrast to mouse Arl5c, human Arl5c is significantly different from the rest paralogs as it does not have a typical G3 box (Supplementary Fig. 4a). Using purified cDNA plasmids as calibration standards, our RT-qPCR revealed that transcripts of Arl5a and b are roughly equal while that of Arl5c is ~ 30 folds less than Arl5a and b in HeLa cells, consistent with the proposal that *Arl5c* gene could be a pseudogene(Rosa-Ferreira et al., 2015). In agreement with their high sequence identity, immobilized GST-Arl5a, Arl5b and mouse Arl5c pulled down Lamtor1-GFP, though Arl5b appeared to retain the most (Supplementary Fig. 4b). These results suggest that Arl5a, Arl5b and mouse Arl5c could have redundant cellular functions but Arl5b possibly contributes most to Ragulator interaction.

### Arl5b colocalizes with Lamtor1 at the endosome and lysosome

When transiently expressed in HeLa cells, C-terminally GFP-tagged wt, QL and TN mutant forms of Arl5a, Arl5b or mouse Arl5c (Fig. 5a; Supplementary Fig. 5a,b) localized to the Golgi, although TN form had greatly reduced Golgi localization with concomitantly increased cytosolic pool. In contrast, human Arl5c-wt-GFP did not localize to the Golgi (Supplementary Fig. 5c). We raised Arl5b-specific polyclonal antibody (Supplementary Fig. 5d-f) and the staining of endogenous Arl5b further confirmed its Golgi localization (Fig. 5b). Similar to Arl1(Lu et al., 2001), the N-terminal myristoylation of Arl5b at Gly of position 2 seemed to be essential for its Golgi localization (Supplementary Fig. 5g). Taking advantage of GLIM (Golgi protein localization by imaging centers of mass), our newly developed quantitative localization method for Golgi proteins (Tie et al., 2016), localization quotients (LQs) of GFP-tagged Arl5a and b were measured to be 0.99 ± 0.02 (n=139) and 0.91 ± 0.02 (n=94), respectively, indicating that they are mainly localized to the *trans*-Golgi.

**Figure 5.**
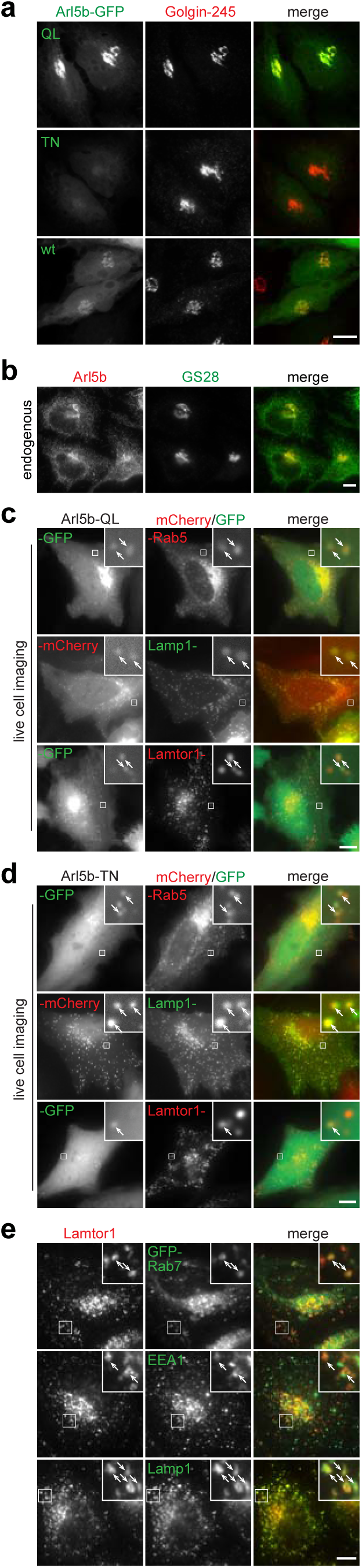
Arl5 localizes to the Golgi, endosome and lysosome. All cells are HeLa cells. (**a**) The Golgi localization of different mutant forms of Arl5b. Cells transiently expressing Arl5b-GFP in QL, TN or wt form were fixed and endogenous Golgin-245 was stained. (**b**) Endogenous Arl5b localizes to the Golgi. HeLa cells were fixed and endogenous Arl5b and GS28 were co-stained. (**c,d**) Arl5b colocalizes with Lamtor1 at the endosome and lysosome. HeLa cells transiently co-expressing indicated GFP or mCherry-tagged proteins were imaged under live cell condition. (**e**) Lamtor1 localizes to the EE, LE and lysosome. Endogenous Lamtor1 was co-stained with exogenously expressed GFP-Rab7, endogenous EEA1 or Lamp1, respectively. In **c-e**, the boxed region was enlarged in the upper right corner to show the colocalization at puncta (denoted by arrows). Scale bar, 10 μm.

Interestingly, in addition to the Golgi pool observed in fixed cells, live-cell imaging revealed that C-terminally mCherry or GFP-tagged Arl5b-QL and -TN also localized to peripheral puncta positive for mCherry-Rab5 (an EE marker) and Lamp1-GFP (a LE or lysosome marker) (Fig. 5c,d), demonstrating the endosomal and lysosomal localization of Arl5b. Most importantly, the peripheral puncta of GFP-tagged Arl5b-QL or TN colocalized extensively with Lamtor1-mCherry (Fig. 5c,d). In contrast, no peripheral puncta were observed in Arl5b-wt-GFP expressing cells under identical condition (Supplementary Fig. 5h). Besides the LE and lysosomal localization of Lamtor1(Nada et al., 2009; Sancak et al., 2010), a substantial amount of Lamtor1 also colocalized with EEA1 (an EE marker) (Fig. 5e). However, Lamtor1 did not localize to the Golgi (Supplementary Fig. 5i). Together with our biochemical data, our data suggest that the interaction between Arl5b and Ragulator can take place on the surface of the endosome and lysosome.

### Arl5b and its effector, GARP, are essential for the AA-stimulated endosome-to-Golgi trafficking

Our findings prompted us to test the hypothesis that Arl5b participates in the AA-stimulated endosome-to-Golgi trafficking. Due to the potential redundancy, endogenous Arl5a, b and c were simultaneously depleted by a mixture of three siRNAs targeting the three paralogs (Fig. 3h; Supplementary Fig. 6a,b). The simultaneous depletion of Arl5a and b significantly blunted the AA-stimulated Golgi trafficking of CD8a-furin (Fig. 3i). Singly depletion of either Arl5a or Arl5b using alternative shRNAs resulted in similar inhibitory effect (Supplementary Fig. 6c-f). GARP complex has recently been identified as the effector of Arl5(Rosa-Ferreira et al., 2015). It localizes to both the TGN and endosomes(Quenneville et al., 2006; Schindler et al., 2015) and functions as a tethering factor in the endosome-to-Golgi trafficking(Bonifacino and Hierro, 2011). There are four subunits in GARP complex: Vps51-54(Bonifacino and Hierro, 2011). Upon depleting endogenous Vps51 or Vps54 (Fig. 3j), the AA-stimulated Golgi trafficking was found substantially attenuated (Fig. 3k). Together, our data demonstrate that Arl5 and its effector, GARP, are essential for the AA-stimulated endosome-to-Golgi trafficking.

**Figure 6.**
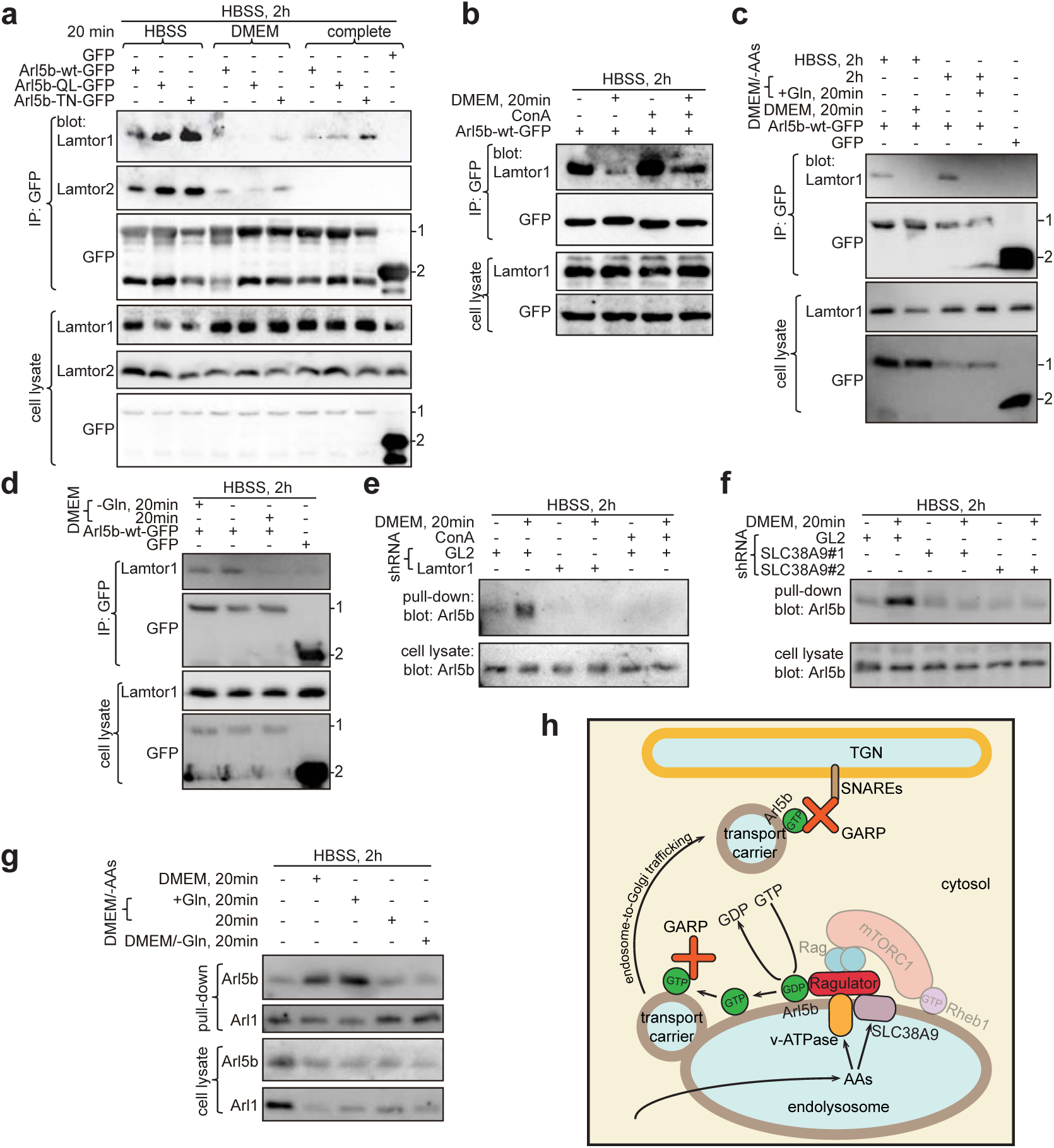
Under AA-sufficiency, Ragulator functions as a GEF for Arl5b downstream of v-ATPase and SLC38A9. (**a**) AAs disrupt Arl5b-Ragulator interaction, regardless of guanine nucleotide binding status of Arl5b. HEK293T cells expressing indicated GFP-fusions were starved in HBSS for 2 h before treatment by indicated medium for 20 min. The resulting cell lysates were incubated with anti-GFP pAb and co-IPs were analyzed by immuno-blotting indicated proteins. (**b**) AA-induced disruption of Arl5b-Ragulator binding is inhibited by conA. HEK293T cells expressing Arl5b-wt-GFP were treated by HBSS for 2 h. Cells were lysed directly or after 20 min treatment of DMEM. For conA treatment, 2.5 μM conA was used throughout the incubation. Co-IPs and immuno-blotting were performed as in **a**. (**c**) Gln is sufficient to disrupt Arl5b-Ragulator binding. HEK293T cells expressing Arl5b-wt-GFP or GFP were starved in HBSS or DMEM/-AAs for 2 h and subsequently incubated with DMEM or DMEM/-AAs supplemented with Gln. Co-IPs and immuno-blotting were performed as in **a**. (**d**) Gln is necessary to disrupt Arl5b-Ragulator binding. HEK293T cells expressing Arl5b-wt-GFP or GFP were starved in HBSS for 2 h. Cells were either directly lysed or after 20 min treatment of indicated medium. Co-IPs and immuno-blotting were performed as in **a**. (**e,f**) v-ATPase, Ragulator and SLC38A9 are essential for the AA-stimulated guanine nucleotide exchange of Arl5b. HeLa cells were subjected to shRNA-mediated knockdown of indicated proteins. Nutrient treatment was performed as in **d**. ConA treatment was performed as in **b**. The resulting cell lysates were incubated with GTP-agarose and pull-downs were analyzed by immuno-blotting endogenous Arl5b. GL2 is a non-targeting control shRNA. (**g**) Gln is necessary and sufficient for the AA-stimulated guanine nucleotide exchange of Arl5b. Nutrient treatment and pull-down were performed as in **e** and **f**. In **a**, **c** and **d**, “1” and “2” indicate Arl5b-(wt, QL or TN)-GFP or GFP band, respectively. (**h**) A working model on how Arl5b integrates the AA-sufficiency signal and regulates the endosome-to-Golgi trafficking. See discussion for details.

### AAs regulate Arl5b-Ragulator interaction

The interaction between heterodimeric Rag GTPases and components of mTORC1 signaling, including mTORC1, Ragulator, v-ATPase, SLC38A9 and folliculin-FNIP1, is regulated by AAs — it becomes strengthened and weakened by AA-starvation and sufficiency, respectively(Bar-Peled et al., 2012; Jung et al., 2015; Petit et al., 2013; Rebsamen et al., 2015; Tsun et al., 2013; Wang et al., 2015; Zoncu et al., 2011). We hypothesized that the binding between Arl5b and Ragulator can also be regulated by AAs. Indeed, compared with AA-starvation (HBSS treatment), a 20 min stimulation by either AAs or a combination of AAs and serum substantially decreased the amount of Lamtor1 and Lamtor2 co-IPed by Arl5b-GFP (Fig. 6a).

Given our observation that Arl5b-QL interacts with Ragulator more weakly than TN, we initially thought that the reduced Arl5b-Ragulator binding under AA-sufficiency could be due to the guanine nucleotide exchange of Arl5b from GDP to GTP form. Indeed, we later found that AA-sufficiency induces the guanine nucleotide exchange of Arl5b (see below). However, when Arl5b-bound guanine nucleotide was locked by using either QL or TN mutation, reduced-binding under AA-sufficiency was still observed for both mutant forms (Fig. 6a). Therefore, the reduction in Arl5b-Ragulator interaction might be resulted from the structural change in Ragulator upon integrating upstream AA-sufficiency signal from v-ATPase and SLC38A9. Consistent with this view, conA treatment increased Arl5b-Ragulator binding during AA-sufficiency (Fig. 6b). Thus, AAs modulate Ragulator’s engagement with Arl5b, similar to previously reported Ragulator-Rag GTPase interaction(Bar-Peled et al., 2012).

Since Gln has the most acute effect on the endosome-to-Golgi trafficking among all AAs, we asked if Gln also plays a significant role in regulating the Arl5b-Ragulator interaction. Following AA-starvation (HBSS or DMEM/-AAs treatment), we observed that Arl5b-Lamtor1 interaction was abolished when cells were subsequently treated with Gln alone (Fig. 6c); in contrast, the interaction remained as strong as that in AA-starvation when cells were treated with DMEM/-Gln (Fig. 6d). Therefore, Gln is sufficient and necessary to disrupt Arl5b-Ragulator interaction.

### Ragulator probably functions as a GEF for Arl5b

Most small GTPases interact with their effectors in GTP-bound form. Hence it seems unusual that Ragulator preferentially interacts with GDP-bound form of Arl5b. Furthermore, we found that Ragulator exhibited higher binding affinity toward guanine nucleotide free Arl5b, which was prepared by EDTA treatment, than either GDP or GTP-bound form of Arl5b (Fig. 4g). Therefore, the role of Ragulator for Arl5b is more consistent with a GEF than an effector. In fact, Ragulator was previously demonstrated to be an AA-regulated GEF for small GTPases, RagA and RagB(Bar-Peled et al., 2012). Our findings thus prompted us to test if Ragulator can similarly function as a GEF for Arl5b. The GEF activity was assayed by GTP-agarose binding assay, in which guanine nucleotide exchange couples Arl5b to agarose-linked GTP and therefore the amount of Arl5b pulled down can be used to measure the GEF activity. Supporting our hypothesis, in control knockdown cells, GTP-agarose pulled down substantially more endogenous Arl5b under AA-sufficiency than starvation (Fig. 6e); the AA-stimulated GTP-binding of Arl5b was blocked by the knockdown of endogenous Lamtor1 or SLC38A9 or conA-mediated inhibition of v-ATPase (Fig. 6e,f). We demonstrated that Gln alone increased while DMEM/-Gln decreased the GTP-binding of Arl5b (Fig. 6g), suggesting that Gln is necessary and sufficient for promoting the GEF activity of Ragulator toward Arl5b. Our data therefore strongly support our hypothesis that Ragulator functions as a GEF for Arl5b by integrating AA-sufficiency signal from v-ATPase and SLC38A9.

## Discussion

Despite our advancement in understanding trafficking processes among organelles, there is a lack of knowledge on how intracellular membrane trafficking processes are regulated in response to extracellular signals. We present here a novel discovery that extracellular AAs, but not growth factors and glucose, regulate the endosome-to-Golgi trafficking in mammalian cells. Under AA-starvation, cargos cycling between the PM and Golgi are arrested in the endosome and the subsequent AA-stimulation rapidly promotes the retrograde trafficking of cargos to the Golgi.

The AA-stimulated endosome-to-Golgi trafficking can affect the cellular metabolism in at least two-folds. First, it is known that the endosome-to-Golgi trafficking prolongs half-lives of post-Golgi cycling cargos by diverting them away from lysosomal degradation pathway. Under AA-sufficiency, longer half-lives of proteins and lipids probably contribute to anabolic processes for cell growth and proliferation. Second, cells can also utilize the AA-stimulated retrograde trafficking to quantitatively adjust certain transporters and receptors on the PM to ensure an optimal uptake of nutrients and engagement with their environment. During AA-starvation, compromised endosome-to-Golgi trafficking can retain more proteins on the PM to acquire more extracellular AAs, as exemplified by yeast Gap1p(Chen and Kaiser, 2002; Roberg et al., 1997). We observed that Gln, one of the most effective nitrogen sources for yeast, most acutely stimulates the endosome-to-Golgi pathway. Gln has the highest concentration in both blood plasma (0.5 - 0.8 mM) and cell culture media (2 - 4 mM). It fuels the tricarboxylic acid cycle and contributes to the biosynthesis of macromolecules; furthermore, it facilitates the uptake of essential AAs and activates mTORC1 signaling pathway(Jin et al., 2016; Wise and Thompson, 2010). In fact many cancer cells heavily rely on Gln for their growth and survival, the complete mechanism of which remains to be elucidated. It is possible that the maintenance of the endocytic retrograde trafficking contributes to the cellular demand for Gln.

We elucidated a signaling pathway from the sensing of AAs to the trafficking of membrane carriers from the endosome to the Golgi. Our data demonstrate that components in the AA-sensing module of mTORC1 pathway, excluding mTORC1 and Rag GTPases, play similar roles in AA-stimulated endosome-to-Golgi trafficking. Hence the AA-regulated retrograde trafficking and mTORC1 signaling share common components, including SLC38A9, v-ATPase and Ragulator, but the two pathways diverge after Ragulator: while the activation of mTORC1 is through Rag GTPases, the promotion of the retrograde trafficking requires the Arl-subfamily small GTPase — Arl5b.

The Golgi-localized Arl5b has been previously known to participate in the endosome-to-Golgi trafficking by interacting with the tethering complex, GARP(Houghton et al., 2012; Rosa-Ferreira et al., 2015). We further discovered that Arl5b and GARP are essential players for the AA-stimulated endosome-to-Golgi trafficking. From our biochemical and imaging data, Ragulator was found to interact with Arl5b on the endolysosome via its scaffolding subunit — Lamtor1. We found that AAs stimulate the guanine nucleotide exchange of Arl5b from GDP to GTP in a Ragulator, v-ATPase and SLC38A9 dependent manner. Our GTP-agarose pull-down data collectively suggest that Ragulator functions as a GEF for the activation of Arl5b, therefore providing a mechanistic connection between endocytic trafficking and the nutrient signaling pathway. We propose here a working model to summarize the AA-regulated signaling pathway that leads to the retrograde trafficking (Fig. 6h). Similar to mTORC1 signaling pathway, the luminal AAs of the endolysosome are first sensed by SLC38A9 and v-ATPase, which in turn signal to Ragulator; activated Ragulator subsequently acts as a GEF to activate Arl5b by guanine nucleotide exchanging; at last, GARP is recruited to the membrane carrier by activated Arl5b and facilitates the tethering and fusion of the budded membrane carriers with the TGN membrane.

We have shown in this study that, in addition to Rag GTPases, Ragulator also possesses GEF activity towards Arl5b on the endolysosomal membrane. It is unknown at this moment if endolysosome-localized Ragulator can contribute to the activation of Arl5b on the Golgi membrane. In agreement with its GEF activity, four subunits of Ragulator, Lamtor2-4, have roadblock domains, which are expected to have GTPase-binding activity(Levine et al., 2013). It is tempting to speculate that, besides Rag and Arl5b, Ragulator might serve as a GEF for other small GTPases on endolysosomal membrane. These small GTPases might respond similarly to AAs via the same sensing mechanism to regulate endolysosomal activities. Thus Ragulator is probably a key node in the AA-regulated signaling network. The endosome-to-Golgi trafficking has been established to regulate physiological and pathological processes such as metazoan development, bacterial toxin invasion, cellular homeostasis and neurological diseases(Burd, 2011; Lu and Hong, 2014). Our discovery therefore implies that nutrient might modulate and affect these processes, an interesting prediction awaiting further study.

## Materials and methods

### Antibodies and small molecules

The following antibodies were purchased from Cell Signaling Technology: rabbit anti-p70S6K polyclonal antibody (pAb) (#9202), rabbit anti-RagA monoclonal antibody (mAb) (#4357), rabbit anti-Lamtor1 mAb (#8975), rabbit anti-Lamtor2 mAb (#8145), rabbit anti-Lamtor3 mAb (#8168) and rabbit anti-Lamtor4 mAb (#13140). The following antibodies were from Abcam: rabbit anti-α-tubulin pAb (#ab4074), rabbit anti-EEA1 pAb (#ab2900), rabbit anti-giantin pAb (#ab24586), rabbit anti-GM130 pAb (#ab52649) and HRP-conjugated protein A (#ab7456). The following antibodies were from Santa Cruz: rabbit anti-GAPDH pAb (#sc-25778), mouse anti-Lamp1 mAb (#sc-20011), mouse anti-GFP mAb (#sc-9996), mouse anti-Myc mAb (#sc-40), mouse anti-HA mAb (#sc-7396) and rabbit anti-Rab7 pAb (#sc-10767). Mouse anti-Lamp1 mAb (H4A3) and mouse anti-CD8a mAb (OKT8) were from Developmental Studies Hybridoma Bank. Rabbit anti-furin pAb (#PA1062) and mouse anti-M6PR mAb (#MA1066) were from Thermo Scientific. Mouse anti-Flag mAb was from Sigma-Aldrich (#F1804). HRP-conjugated goat anti-mouse (#176516) and anti-rabbit IgG antibodies (#176515) were from Bio-Rad. Mouse anti-GM130 mAb (#610823), mouse anti-EEA1 mAb (#610456) and mouse anti-syntaxin6 mAb (#51-9002100) were from BD Biosciences. Rabbit anti-RUFY1 pAb (#13498-1-AP) was from Proteintech. Alexa Fluor conjugated goat anti-mouse and anti-rabbit IgG antibodies were from Invitrogen. Rabbit anti-Arl5b antibody is generated as described below. Rabbit anti-Arl1 pAb was previously described(Lu and Hong, 2003).

The following small molecule inhibitors are commercially available: Concanamycin A (Abcam, #ab144227); Torin 1(Tocris Bioscience, #4247); rapamycin (InvivoGen, #tlrl-Rap). GMPPNP (#G0635) and GDP (#G7127) were from Sigma-Aldrich.

### Yeast-two hybrid screening

The yeast-two hybrid screening was conducted using Arl5b-QL in pGBKT7 as a bait to screen human kidney cDNA library pre-transformed yeast (Clontech), similar to previously described(Lu et al., 2001).

### Cell culture, transfection, surface-labeling and acid wash

HeLa, BSC-1, HEK293T and 293FT cells were maintained in high glucose DMEM (GE Healthcare Life Science) supplemented with 10% fetal bovine serum (FBS) (Gibco) at 37°C in 5% CO2 incubator. HeLa, BSC-1 and HEK293T cells were transfected using polyethylenimine (Polysciences, Inc.). Transfection was performed when cells reached 70-80% confluency according to standard protocol.

DMEM-base was prepared using 100x MEM vitamin solution (Invitrogen, #11120052), inorganic salts, glucose and sodium pyruvate according to the formulation of DMEM from Invitrogen (#11965) leaving out all AAs. Selective AA(s) was(were) added to DMEM-base to make corresponding media containing defined AAs. DMEM/-Gln and DMEM/-Leu were prepared by supplying Leu and Gln, respectively, to DMEM/-Gln/-Leu (MP Biomedicals, #1642149). HBSS was prepared according to the formulation of Invitrogen HBSS (#14025). Except Gln (Invitrogen) and His (Fluka), all AAs were from Sigma-Aldrich. Concentrations of individual AAs in nutrient media were either according to the formulation of DMEM of Invitrogen (#11965) or as indicated in the text. Dialyzed-serum was prepared by dialyzing the FBS in 3.5 kd molecular weight cut-off dialysis tubing (Thermo Fisher, #68035) against PBS followed by passing through a syringe-driven 0.22 μm filter unit (Sartorius).

Surface-labeling was conducted by incubating live cells with anti-CD8a antibody (OKT8) for 1 h on ice. Unbound antibody was subsequently washed away by ice cold PBS and cells were incubated in AA-starvation or -sufficiency medium at 37 °C for certain length of time before being processed for imaging. Acid wash was conducted to strip-off surface-exposed CD8a antibody that binds to CD8a-furin. Briefly, live cells were incubated with ice cold 0.2 M acetic acid in 0.5 M NaCl for 4 min and subsequently washed extensively by ice cold PBS. Cells were then subjected to endocytic trafficking at 37 °C in indicated medium.

### Lentivirus-mediated transduction of shRNAs or fusion proteins in HeLa cells

293FT cells seeded in a 6-well-plate were transfected with shRNA in pLKO.1 vector or a mixture of pLVX expression construct, pLP1, pLP2 and pLP/VSVG using Lipofectamine 2000 (Invitrogen). Cells were incubated at 37 °C for 18 h and replaced with fresh medium to incubate for another 24-48 h to harvest virus-containing supernatant. The supernatant was passed through 0.45 μm filter (Invitrogen) and used immediately. For lentivirus-mediated transduction, HeLa cells were incubated with the virus supernatant in the presence of 8 μg/ml of polybrene (Sigma #H9268) 24-48 h before an assay. For the expression of an exogenous protein after knockdown, transient transfection was conducted 48 h after the addition of lentivirus and cells were further incubated for 24 h before the indicated-assay. To select CD8a-furin stable cells, virus-infected HeLa cells were cultured in the medium containing 1 μg/ml puromycin (Sigma-Aldrich). The pooled stable cells were subjected to limited dilution and cultured in a 96-well plate to expand single cell colonies. The colonies were further screened by immunofluorescence or Western blot.

### RNAi-mediated knockdowns

The following siRNAs were purchased from GE Dharmacon: luciferase GL2 (#D-001100-01-20) and siRNA SMART pools for human Lamtor1 (#L-020916- 02-0005), human Arl5a (#L-012408-00-0005), Arl5b (#L-017861-02-0005) and Arl5c (#L-030887-02-0005). siRNAs were transfected to HeLa cells using Lipofectamine 2000 according to manufacturer’s protocol. For the expression of an exogenous protein after knockdown, transfections were conducted 24 h after siRNA transfection. 48 h after the transfection of siRNA, cells were processed for assays.

### Reverse transcription real-time PCR (RT-qPCR)

Total RNA was purified using Trizol reagent (Invitrogen) from HeLa cells according to standard protocol. Reverse transcription primed by random nonamer primers was conducted using nanoScript 2 Reverse Transcription kits (Primerdesign). The real-time PCR was subsequently performed on a Bio-Rad CFX96 Touch^™^ real-time PCR detection system using SYBR green based PrecisionFAST with LOW ROX qPCR kit (Primerdesign). The specificity of PCR primers were verified by both melt curves and agarose gel electrophoresis. The primer pairs are listed below: Arl5a (5’-GTT AGC GCA TGA GGA CCT AAG-3’, 5’- CTT GGC ACA ATC CCT CGC CAG TTA C-3’), Arl5b (5’-TGG CTC ATG AGG ATT TAC GGA AG-3’, 5’-CCT TGG CAT AAC CCT TCT CCT GTG-3’), hArl5c (5’- TGG CCC ATG AGG CTC TAC AGG ATG-3’, 5’-TCC ATC CAC TGA AGT CTG GCA G-3’), Vps51 (5’-CTC AGC CAC AGA CAC CAT CCG G-3’, 5’-GCG AGC GCT GAA GTC GGT GAT C-3’), Vps54 (5’-GTT GTT GTG AAG CTT GCA GAT CAG-3’, 5’-TGT TGC CTT CAC TCT CTG TAG G-3’), SLC38A9 (5’-CCT AGC ATT TTC CAT GTG CTG-3’, 5’-GCT CCT GAA TAT CTT ATG ATC CCT CC-3’) and Lamtor3 (5’-CCT GTT ATT AAA GTG GCAAAT GAC AAT GC-3’, 5’-TTG AAC CAC CTG GTA GGT GTT ATA G-3’).

### Subcellular fractionation by sucrose gradient ultracentrifugation

HeLa cells stably expressing CD8a-furin were cultured in two Φ15 cm Petri-dishes to 70% confluency and treated with HBSS for 2 h. Cells on one plate were further incubated with DMEM for 20 min. Next, cells on both plates were lysed in 10% sucrose buffer (10% sucrose, 3 mM imidazole pH 7.4, 1 mM EDTA) by repeatedly extruding them through a 25 1/2 gauge needle. The cell lysates were subsequently centrifuged at 1,000 g for 10 min. The resulting two supernatants were separately loaded on the top of two tubes containing 10 - 40% continuous sucrose gradient, which were then subjected to ultracentrifuge in SW28 rotor (Beckman) at 140,000 g and 4 °C for 5 h. After the centrifugation, samples in tubes were collected into 20 fractions with 1.85 ml per fraction. Proteins within each fraction were pelleted down using methanol/chloroform method(Wessel and Flugge, 1984), dissolved in SDS-sample buffer and analyzed by Western blot.

### Purification of GST and His-tagged fusion proteins

This was performed as previously described(Madugula and Lu, 2016; Mahajan et al., 2013; Zhou et al., 2013).

### Generation of anti-Arl5b rabbit polyclonal antibody

His-tagged Arl5b was purified under denaturation condition using 8 M urea as previously described(Mahajan et al., 2013). The denatured protein was used to immunize rabbits and anti-sera were collected by Genemed Synthesis Inc. The affinity purification of Arl5b antibody was conducted using bead-crosslinked GST-Arl5b as previously described(Madugula and Lu, 2016).

### Guanine nucleotide exchange of GST-Arl5b

Glutathione bead-immobilized GST-Arl5b was washed twice with the exchange buffer (20 mM HEPES, pH 7.4, 100 mM NaCl, 10 mM EDTA, 5 mM MgCl2 and 1 mM DTT) and incubated with the buffer supplemented with 10 unit/ml calf intestinal alkaline phosphatase (New England Biolab) at room temperature for 2 h. Next, beads were washed by the exchange buffer and incubated with the same buffer supplemented with 0.5 mM GMPPNP or GDP (final concentration) for 1 h at the room temperature. 5 mM (final concentration) MgCl2 was subsequently added to the system and beads were further incubated for 1 h at the room temperature. The exchanged GST-Arl5b on beads was stored at 4 °C until use.

### Co-Immunoprecipitation and pull-down

HEK293T or HeLa cells were transfected by indicated DNA construct to express the exogenous protein and/or treated with the AA-starvation or -sufficiency medium as indicated in the text. After washing cells with ice cold PBS, cells were lysed in lysis buffer (40 mM HEPES, pH 7.4, 150 mM NaCl, 1% Trition X-100, 2.5 mM MgCl2, 1 mM PMSF) and cleared by centrifugation at 16 kg for 30 min. In co-IPs or pulldowns involving Ragulator, Triton X-100 was substituted by 0.3% CHAPS (Sigma-Aldrich). Cell lysates were subsequently incubated with ~ 1μg antibody against the protein of interest, 15 μl GFP-Trap beads (ChromoTek) or 10-40 μg GST-fusion protein immobilized on glutathione beads for 4-14 h in a cold room. When antibody was used, the antigen-antibody complex was subsequently captured by 15 μl pre-washed Protein A/G beads (Pierce). After washing beads extensively with the lysis buffer, bound proteins were eluted by boiling in SDS-sample buffer and resolved in 8-12% SDS-PAGE. Western blot was subsequently conducted to detect bound proteins according to standard protocol. Separated proteins were transferred to polyvinyl difluoride membrane (Bio-Rad). After primary and HRP-conjugated secondary antibody incubation, the chemiluminescence signal was detected by a cooled charge-coupled device camera (LAS-4000, GE Healthcare Life Sciences).

### GTP-agarose pull-down

HeLa cells suspended in binding buffer (20 mM HEPES, pH 8.0, 150 mM NaCl, 10 mM MgCl2, EDTA-free protease inhibitor (Roche)) were lysed by repeatedly extruding them through a 25 1/2 gauge needle. Lysates were cleared by centrifugation at 16 kg for 30 min and then subjected to incubation with 100 μl GTP-agarose beads (bioWORLD) for 1 h at 4 °C. Beads were washed for three times with binding buffer and bound proteins were eluted by boiling in SDS-sample buffer and analyzed by Western blot.

### Immunofluorescence labeling

It is performed as previously described(Madugula and Lu, 2016).

### Fluorescence microscopy

High resolution images were acquired under an inverted wide-field microscope system, comprising Olympus IX83 equipped with a Plan Apo oil objective lens (63X or 100X, NA 1.40), a motorized stage, motorized filter cubes, a scientific complementary metal-oxide semiconductor camera (Neo; Andor) and a 200 W metal-halide excitation light source (Lumen Pro 200; Prior Scientific,). Dichroic mirrors and filters in filter turrets were optimized for GFP/Alexa Fluor 488, mCherry/Alexa Fluor 594 and Alexa Fluor 647. The microscope system was controlled by MetaMorph software (Molecular Devices) and only center quadrant of the camera sensor was used for imaging.

### Image quantification

Image analysis was performed in ImageJ (http://imagej.nih.gov/ij/). In transient transfection, cells have various levels of expression of the reporter. Therefore, different cells should have distinct background fluorescence intensities. The region of interest (ROI) of the cell was manually drawn by tracing the cell’s contour. The ROI of the Golgi was generated by intensity thresholding using the co-stained endogenous giantin signal. The image is background-subtracted by using ROIs outside cells. In the channel of the reporter fluorescence, A_cell_ and A_Golgi_ are the area (in pixels) of the cell and the Golgi ROI respectively, while I_cell_ and I_Golgi_ are the mean intensity of the cell and the Golgi ROI respectively. f is a constant value between 0 and 1. f=0.5 was used for image quantification with either transfected or endogenous reporters. The fraction of Golgi-localized reporter was calculated as (I_Golgi_-f*I_cell_)*A_Golgi_/((1-f)*I_cell_*A_cell_). Other image analysis was conducted as described in the text. In each image to be quantified, all cells positive for the reporter of interest were analyzed.

### Statistics

*P*-values were determined using Student’s *t*-test (unpaired and two-tailed) in Excel (Microsoft).

## Acknowledgement

We would like to thank W. Hong lab (Institute of Molecular and Cell Biology, Singapore) for the initial support of this work and D. Sabatini, T. Kirchhausen and D. Root for sharing DNA plasmids. This work was supported by the following grants to L.L.: NMRC/CBRG/007/2012, MOE AcRF Tier1 RG132/15, Tier1 RG48/13 and Tier2 MOE2015-T2-2-073.

## Author contribution

L.L. conceived, designed and supervised the study. M.S., B.C., B.K.B., Y.Z., D.M. and H.C.T. performed experiments. M.S., B.C. and L.L. analyzed data. L.L. wrote the manuscript.

## Supplementary Information

### DNA plasmids

#### Constructs of Arl5

To construct <u>Arl5a-GFP</u>, the coding sequence (CDS) of human Arl5a was PCR amplified from a cDNA clone (GenBank Accession No.: NM_012097) using a pair of oligonucleotides (5’-CCG GAA TTC GCC ACC ATG GGA ATT CTC TTC ACT AGA ATA-3’ and 5’-CGC GGA TCC CGT CTA ATC TTA AGT CGT GAC ATC-3’) as primers and ligated into EcoRI/BamHI digested pEGFP-N1 vector (Clontech) using the same sites. To construct <u>Arl5a-Q70L-GFP</u>, two PCR amplifications were performed using Arl5a-GFP as the template and primer pairs (5’-CCG GAA TTC GCC ACC ATG GGA ATT CTC TTC ACT AGA ATA-3’ and 5’-AGA ACG AAG AGA TTC AAG GCC ACC AAT ATC CCA-3’) and (5’-TGG GAT ATT GGT GGC CTT GAA TCT CTT CGT TCT-3’ and 5’-CGC GGA TCC CGT CTA ATC TTA AGT CGT GAC ATC-3’). The two PCR fragments were mixed and subjected to a second round of PCR amplification using the first and the fourth primer. The resulting PCR product was digested by EcoRI/BamHI and ligated into pEGFP-N1 vector using the same sites. <u>Arl5a-T30N-GFP</u> was similarly constructed using primer pairs (5’-CCG GAA TTC GCC ACC ATG GGA ATT CTC TTC ACT AGA ATA-3’ and 5’-TTG GTA AAG AAT GGT AGT TTT CCC TGC ATT ATC-3’) and (5’-GAT AAT GCA GGG AAA ACT ACC ATT CTT TAC CAA-3’ and 5’-CGC GGA TCC CGT CTA ATC TTA AGT CGT GAC ATC-3’). To construct <u>GST-Arl5a</u>, Arl5a-GFP was digested by EcoRI/BamHI and the released insert was ligated into pGEB vector^1^ using the same sites.

To construct <u>Arl5b-GFP</u>, the CDS of human Arl5b was PCR amplified from a cDNA clone (IMAGE clone: 5782580, GenBank Accession No: BQ270027) using a pair of oligonucleotides (5’-CCG GAA TTC GCC ACC ATG GGG CTG ATC TTC GCC AAA CTG TG-3’ and 5’-CTA GCT GGA TCC CGT CTC ACA CCA ATC CGG GAG-3’) as primers and ligated into EcoRI/BamHI digested pEGFP-N1 vector using the same sites. To construct <u>Arl5b-QL-GFP</u>, two PCR amplifications were performed using Arl5b-GFP as the template and primer pairs (5’-CCG GAA TTC GCC ACC ATG GGG CTG ATC TTC GCC AAA CTG TG-3’ and 5’-GAT CGC AGA GAC TCA AGA CCA CCA ATA TCC CAC-3’) and (5’-GTG GGA TAT TGG TGG TCT TGA GTC TCT GCG ATC-3’ and 5’-CTA GCT GGA TCC CGT CTC ACA CCA ATC CGG GAG-3’). The two PCR fragments were mixed and subjected to a second round of PCR amplification using the first and the fourth primer. The resulting PCR product was then digested by EcoRI/BamHI and ligated into pEGFP-N1 vector using the same sites. <u>Arl5b-TN-GFP</u> was similarly constructed using primer pairs (5’-CCG GAA TTC GCC ACC ATG GGG CTG ATC TTC GCC AAA CTG TG-3’ and 5’-GAT AAT GCA GGG AAA AAT ACC ATT CTT TAC C-3’) and (5’-GGT AAA GAA TGG TAT TTT TCC CTG CAT TAT C-3’ and 5’-CTA GCT GGA TCC CGT CTC ACA CCA ATC CGG GAG-3’). To construct <u>Arl5b-QL or TN-mCherry</u>, <u>Arl5b-QL or TN-GFP</u> was digested with EcoRI/BamHI and the insert released was ligated into pmCherry-N1 using the same sites. To construct <u>Arl5b-His</u>, the CDS of Arl5b was PCR amplified using a pair of oligonucleotides (5’-ACG ATA AGA TCT GCC ACC ATG GGG CTG ATC TTC GCC AAA C-3’ and 5’-AGT TCA AAG CTT TCT CAC ACC AAT CCG GGA GGT CAT CCA C-3’) as primers and ligated into BglII/HindIII digested pET-30a vector (Novagen) using the same sites. To construct <u>GST-Arl5b-wt, QL or TN</u>, Arl5b-wt, QL or TN-GFP was digested with EcoRI/BamHI and the resulting fragment was ligated into pGEB vector using the same sites. To construct <u>Arl5b-G2A-QL-GFP</u>, PCR amplification was performed using Arl5b-QL-GFP as the template and a pair of oligonucleotides (5’-ACC GCA GAA TTC GCC ACC ATG GCG CTG ATC TTC GCC AAA CTG-3’ and 5’-CAT GAC GGA TCC CGT CTC ACA CCA ATC CGG GAG GTC ATC-3’) as primers. The resulting PCR product was digested by EcoRI/BamHI and ligated into pEGFP-N1 vector using the same sites. To construct <u>Arl5b-QL in pGBKT7</u> vector, Arl5b-QL-GFP was digested by EcoRI/BamHI and the released insert was ligated into pGBKT7 vector (Clontech) using the same sites.

To construct human and mouse <u>Arl5c-GFP</u>, the CDS of human and mouse Arl5c were amplified from two cDNA clones (GenBank Accession No.: NM_001143968 and BC065791.1, respectively) using primer pairs (5’-GCA CCG GAA TTC GCC ACC ATG GGA CAG CTG ATC GCC-3’ and 5’-CTA GCT GGA TCC CGG TTA GCA GCG GCC TGA G-3’) and (5’-GCG ATC GAA TTC GCC ACC ATG GGA CAG CTG ATA GCC AAG-3’ and 5’-CAC TAC GGA TCC CCG TTG GCG GTG GCC TGA GCT TGC AT-3’), respectively. The resulting PCR products were digested with EcoRI/BamHI and inserted into same enzymes digested pEGFP-N1 vector. To construct <u>mArl5c-QL-GFP</u>, two PCR amplifications were performed using mouse Arl5c-GFP as the template and primer pairs (5’-GCG ATC GAA TTC GCC ACC ATG GGA CAG CTG ATA GCC AAG-3’ and 5’-GCC TCC AGG CCC CCT AGG TCC CAC ATG-3’) and (5’-CAT GTG GGA CCT AGG GGG CCT GGA GGC-3’ and 5’-CTA GCT GGA TCC CGT CTC ACA CCA ATC CGG GAG-3’). The two PCR fragments were mixed and subjected to a second round of PCR amplification using the first and the fourth primer. The resulting PCR product was digested by EcoRI/BamHI and ligated into pEGFP-N1 vector using the same sites. <u>mArl5c-TN-GFP</u>, was similarly constructed using primer pairs (5’-GCG ATC GAA TTC GCC ACC ATG GGA CAG CTG ATA GCC AAG-3’ and 5’-GAG AAT GGT GTT CTT CCC TGC-3’) and (5’-GCA GGG AAG AAC ACC ATT CTC-3’ and 5’-CTA GCT GGA TCC CGT CTC ACA CCA ATC CGG GAG-3’). To construct GST-mArl5c, mArl5c-GFP was digested by EcoRI/BamHI and the released insert was ligated into pGEB vector using the same sites.

#### CD8a chimeras

CD8a-fused furin, CI-M6PR, CD-M6PR and sortilin in pCI-neo vector (Promega) were previously described^2^.

#### Constructs of Lamtors

To construct <u>Lamtor1-GFP</u>, the CDS of Lamtor1 was PCR amplified using a full length clone, which was recovered from our yeast-two hybrid screening, as the template and a pair of oligonucleotides (5’-GAC TAG CTC GAG ATG GGG TGC TGC TAC AGC AGC-3’ and 5’-GAA CTC GAA TTC GTG GGA TCC CAA ACT GTA CAA CCA G-3’) as the primer pair. The resulting PCR product was digested by XhoI/EcoRI and ligated into pEGFP-N1 vector using the same sites. To construct <u>Lamtor1-mCherry</u>, Lamtor1-GFP was digested by XhoI/EcoRI and the insert released was ligated into pmCherry-N1 using the same sites. To construct <u>Lamtor1-Myc</u>, oligonucleotides (5’-AAT TCA GTA CTC AGA ACA AAA ACT CAT CTC AGA AGA GGA TCT GTA AAG C-3’ and 5’-GGC CGC TTT ACA GAT CCT CTT CTG AGA TGA GTT TTT GTT CTG AGT ACT G-3’) were annealed to generate a fragment encoding Myc-tag and ligated into EcoRI/NotI digested Lamtor1-GFP using the same sites. <u>Lamtor1-Flag</u> was similarly constructed by ligating annealed oligonucleotides (5’-GAT CCA GTA CTC GAC TAC AAA GAC GAT GAC GAC AAG TAA AGC-3’ and 5’-GGC CGC TTT ACT TGT CGT CAT CGT CTT TGT AGT CGA GTA CTG-3’) to EcoRI/NotI digested Lamtor1-GFP.

To construct <u>GFP-Lamtor1(1-20</u>) and <u>GFP-Lamtor1(1-39)</u>, CDSs of 1-20 and 1-39 AAs of Lamtor1 were PCR amplified using Lamtor1-GFP as the template and primer pairs (5’-CAG ATC CGC TAG CGC TAC CGG TCG CCA CCA TGG TG-3’ and 5’-GAA CTC GAA TTC GTC ACT TCC GCT CCT CTC GGT CCT GG-3’) and (5’-CAG ATC CGC TAG CGC TAC CGG TCG CCA CCA TGG TG-3’ and 5’-GAA CTC GAA TTC GTC AGT TGG GCT CGG CTC CAT TGA GAG C-3’), respectively. The resulting fragments were digested by NheI/EcoRI and ligated into pEGFP-C3 vector (Clontech) using the same restriction sites. Similarly, <u>GFP-Lamtor1(1-81)</u> and <u>GFP-Lamtor1(1-121)</u> were constructed by primer pairs (5’-GAA CTC GAA TTC GTC AGT ACT CAT GCT GCT CCA TGC CCT G-3’and 5’-GAC TAG CTC GAG ATG GGG TGC TGC TAC AGC AGC-3’) and (5’-GAA CTC GAA TTC GTC AAC TGG CCA GCA CTT GGT GGG GC-3’ and 5’-GAC TAG CTC GAG ATG GGG TGC TGC TAC AGC AGC-3’), respectively, using XhoI/EcoRI sites.

To construct <u>Lamtor1-G2A-Strep-Myc</u>, which contains G2A mutation, a Strep-tag and Myc tag, the CDS of Lamtor1 was PCR amplified using Lamtor1-GFP as the template and primers (5’-CTA GTC CTC GAG ATG GCA TGC TGC TAC AGC A-3’ and 5’-GTC ACT GTC GAC TGT TTT TCG AAC TGC GGG TGG CTC CAC GAT CCA CCT CCC GAT CCA CCT CCG GAA CCT CCA CCT TTC TCG AAC TGC GGG TGG CTC CAT GCT GAT GGG ATC CCA AAC TGT ACA AC-3’). The PCR product was digested by XhoI/SalI and ligated into pMyc-N1 vector^3^ using the same restriction sites. To construct <u>Lamtor2-GFP</u>, the CDS of Lamtor2 was PCR amplified from a cDNA clone (GenBank Accession No.: BC024190) using primers (5’-GTA ATG GA ATT CGA GCC ACC ATG CTG CGC CCC AAG GCT TTG-3’ and 5’- CA CTA CGG ATC CAA AGA TGC CGC CAC TTG GGT G-3’). The resulting PCR product was digested by EcoRI/BamHI and ligated into pEGFP-N1 vector using the same sites. To construct <u>Myc-Lamtor2</u>, Lamtor2-GFP was digeste by EcoRI/NotI to release GFP and the resulting vector was ligated with the annealed oligonucleotides (5’-AAT TCA GTA CTC AGA ACA AAA ACT CAT CTC AGA AGA GGA TCT GTA AAG C-3’ and 5’-GGC CGC TTT ACA GAT CCT CTT CTG AGA TGA GTT TTT GTT CTG AGT ACT G-3’). To construct <u>GFP-Lamtor4</u>, the CDS of Lamtor4 was PCR amplified using a cDNA clone (IMAGE clone: 53000314, GenBank Accession No.: BI598677.1) and the following primer pair (5′-GCG ATC GAA TTC ACT TCT GCG CTG ACC CAG GGG CTG-3′ and 5′-GCG ATC GGA TCC TCA GAC ATC AAT GGG CTC CCG ACC-3′). The resulting fragment was digested by EcoRI/BamHI and ligated into pEGFP-C2 using the same restriction sites. <u>Lamtor5-GFP</u> was similarly constructed using a cDNA clone (IMAGE clone: 53000314, GenBank Accession No.: BI598677.1) and the following primer pair (5′-GCG ATC GAA TTC GAG CCA GGT GCA GGT CAC CTC GAC-3′ and 5′-GCG ATC GGA TCC TCA AGA GGC CAT TTT GTG CAC TGC C-3′). To construct <u>DMyc-Lamtor3, DMyc-Lamtor4 and DMyc-Lamtor5</u>, their CDSs were PCR amplified from a cDNA clone (IMAGE clone: 4808855, GenBank Accession No.: BC026245), GFP-Lamtor4 and GFP-Lamtor5 using primer pairs (5’-CTA GTC GAA TTC AAT GGC GGA TGA CCT AAA GCG A-3’ and 5’-CTA CTC GTC GAC TTA AGA AAC TTC CAC AAC TTG TC-3’), (5’-CTA GTC GAA TTC AAT GAC TTC TGC GCT GAC CCA-3’ and 5’-CTA CTC GTC GAC TCA GAC ATC AAT GGG CTC CC-3’) and (5’-CTT GGA GAA TTC AAT GGA GCC AGG TGC AGG TC-3’ and 5’-CTT GTA GTC GAC TCA AGA GGC CAT TTT GTG CAC-3’), respectively. The resulting PCR products were digested by EcoRI/SalI and ligated into the digested pDMyc-neo vector^2^ using the same sites, respectively. To construct <u>DHA-Lamtor3</u>, DMyc-Lamtor3 was digested by EcoRI/SalI and the released insert was subsequently ligated into pDHA-neo vector. Both pDMyc-neo and pDHA-neo vectors have tandem or double tags and the same multiple cloning sites.

#### Lentivirus expression constructs

To construct <u>CD8a-furin in pLVX-puro</u> vector, the fragment encoding CD8a-furin was PCR amplified from CD8a-furin in pCI-neo using oligonucleotides (5’-GTC TAG AAT TCA GCC ACC ATG GCC TTA CCA GTG ACC GCC TTG C-3’ and 5’-GAC CTG TCT AGA TTA GAG GGC GCT CTG GTC TTT GAT AAA GGC G-3’) as primers. The resulting fragment was digested by EcoRI/XbaI and ligated into pLVX-Puro vector (Clontech) using the same sites.

#### shRNA constructs

To construct <u>GL2 shRNA in pLKO.1</u> vector, oligonucleotide (5’-CCG GAA CGT ACG CGG AAT ACT TCG ACT CGA GTC GAA GTA TTC CGC GTA CGT TTT TTT G-3’ and 5’-AAT TCA AAA AAA CGT ACG CGG AAT ACT TCG ACT CGA GTC GAA GTA TTC CGC GTA CGT T-3’) were annealed and ligated into AgeI/EcoRI digested pLKO.1 vector (Addgene # 10878; a gift from D. Root). shRNAs targeting <u>Arl5a, Arl5b, Vps51 #1, Vps51 #2, Vps54 #1, Vps54 #2, SLC38A9 #1</u> <u>and SLC38A9 #2 in pLKO.1</u> vector were similarly constructed using the following oligonucleotide pairs (5’-CCG GAA TGA TCT CTA CTG ACC TCT TCT CGA GAA GAG GTC AGT AGA GAT CAT TTT TTT G-3’ and 5’-AAT TCA AAA AAA TGA TCT CTA CTG ACC TCT TCT CGA GAA GAG GTC AGT AGA GAT CAT T-3’), (5’-CCG GAA TAC CTC ACC CTT AGT TCA ACT CGA GTT GAA CTA AGG GTG AGG TAT TTT TTT G-3’ and 5’-AAT TCA AAA AAA TAC CTC ACC CTT AGT TCA ACT CGA GTT GAA CTA AGG GTG AGG TAT T-3’), (5’-CCG GAA CCT CTT GAG CAA TAT CCA GCT CGA GCT GGA TAT TGC TCA AGA GGT TTT TTT G-3’ and 5’-AAT TCA AAA AAA CCT CTT GAG CAA TAT CCA GCT CGA GCT GGA TAT TGC TCA AGA GGT T-3’), (5’-CCG GAA CGT ATT GAT GTG TTC AGC CCT CGA GGG CTG AAC ACA TCA ATA CGT TTT TTT G-3’ and 5’-AAT TCA AAA AAA CGT ATT GAT GTG TTC AGC CCT CGA GGG CTG AAC ACA TCA ATA CGT T-3’), (5’-CCG GAA CAT TGC TCA CCA GAT CTC TCT CGA GAG AGA TCT GGT GAG CAA TGT TTT TTT G-3’ and 5’-AAT TCA AAA AAA CAT TGC TCA CCA GAT CTC TCT CGA GAG AGA TCT GGT GAG CAA TGT T-3’), (5’-CCG GAA CCA GCT GAA GTT CTT ATT GCT CGA GCA ATA AGA ACT TCA GCT GGT TTT TTT G-3’ and 5’-AAT TCA AAA AAA CCA GCT GAA GTT CTT ATT GCT CGA GCA ATA AGA ACT TCA GCT GGT T-3’), (5’-CCG GGC CTT GAC AAC AGT TCT ATA TCT CGA GAT ATA GAA CTG TTG TCA AGG CTT TTT G-3’ and 5’-AAT TCA AAA AGC CTT GAC AAC AGT TCT ATA TCT CGA GAT ATA GAA CTG TTG TCA AGG C-3’) and (5’-CCG GCC TCT ACT GTT TGG GAC AGT ACT CGA GTA CTG TCC CAA ACA GTA GAG GTT TTT G-3’ and 5’-AAT TCA AAA ACC TCT ACT GTT TGG GAC AGT ACT CGA GTA CTG TCC CAA ACA GTA GAG G-3’).

The following shRNA constructs in pLKO.1 were gifts from D. Sabatini. Lamtor1 (Addgene: #26631), Lamtor3 (Addgene: #26632), RagA #1 (Addgene: #30319), RagB #1 (Addgene: #26627).

#### Other constructs

To construct <u>furin-GFP or mCherry</u>, the full length CDS of furin was PCR amplified using a cDNA clone (GenBank Accession No.: BC012181.1) as the template and the following oligonucleotides (5’-CAG ATC TCG AGC TCA AGC TTC GAA TTC GCC ACC ATG GAG CTG AGG CCC TGG-3’ and 5’-GAT CCC GGG CCC GCG GTA CCG TCG ACC CGA GGG CGC TCT GGT CTT TG-3’) as primers. The PCR fragment was digested by EcoRI/SalI and ligated into pEGFP-N1 or pmCherry-N1 vector, respectively, using the same sites. Myc-SNX3 was previously described^4^. Lamp1-GFP, GFP-Rab7, mCherry-Rab5, TfR-GFP were gifts from T. Kirchhausen.

---

**Supplementary Figure 1.**
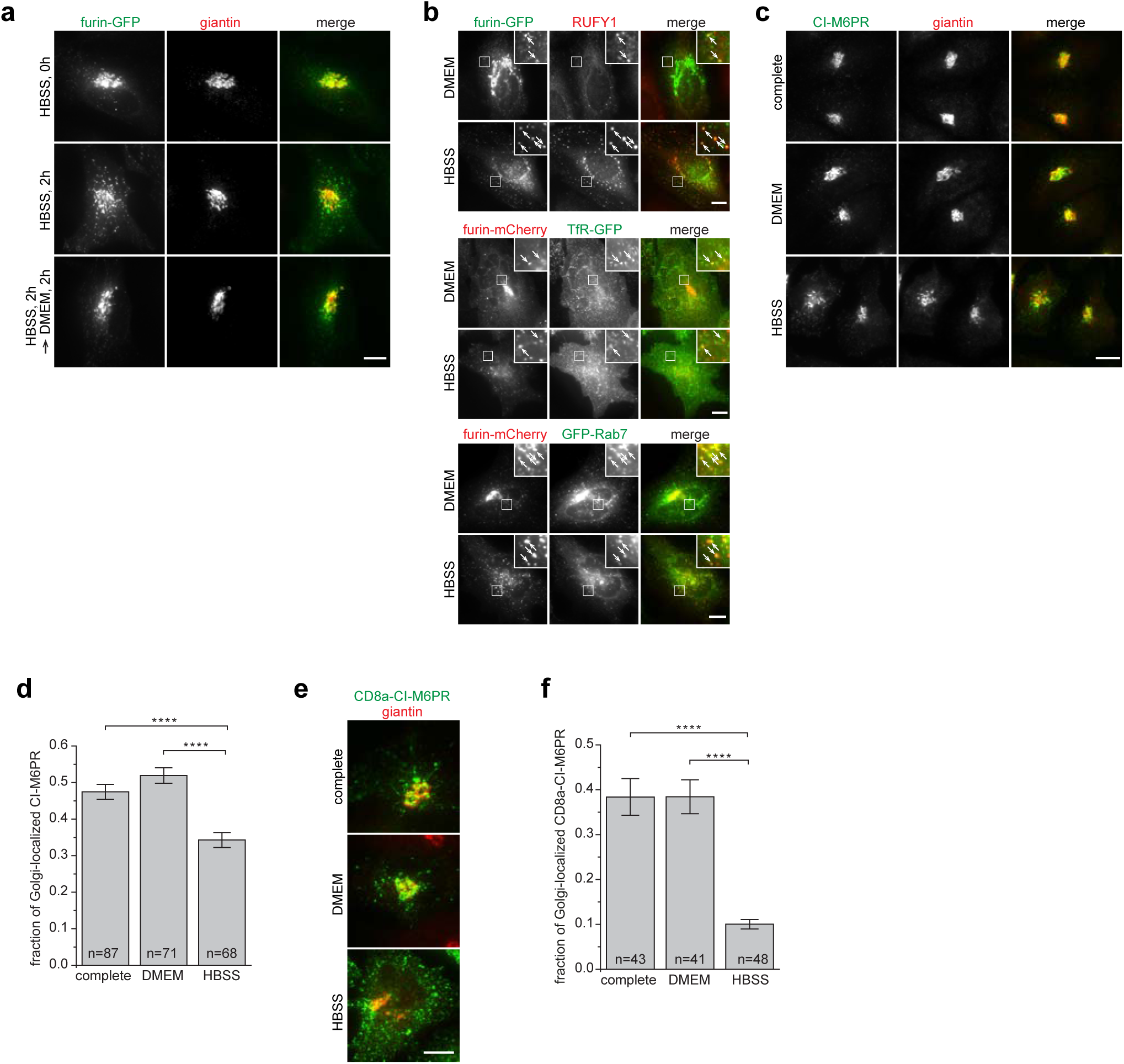
Nutrient starvation induces endosomal translocation of TGN membrane proteins. All cells are HeLa cells. (**a**) The localization of furin-GFP during HBSS and DMEM treatment. Cells were processed as described in Fig. 1d. (**b**) The endosomal localization of furin-GFP or furin-mCherry increases at the expense of its Golgi pool during nutrient starvation. Cells expressing furin-GFP singly (top two panels), furin-mCherry together with GFP-TfR (middle two panels) and furin-mCherry together with GFP-Rab7 (lower two panels) were treated with indicated medium for 2 h. Cells in the top two panels were stained for endogenous RUFY1. Boxed regions are enlarged to demonstrate the colocalization (indicated by arrows). (**c,d**) Nutrient starvation induces significant endosomal translocation of endogenous CI-M6PR. Cells treated with indicated medium for 1 h and endogenous CI-M6PR and giantin were stained. The fraction of Golgi-localized CI-M6PR is quantified in **d**. (**e,f**) Nutrient starvation induces significant endosomal translocation of CD8a-CI-M6PR. The experiment was conducted similarly to **c** and **d** except that cells expressing CD8a-CI-M6PR were used and CD8a was stained. Complete, complete medium; n, the number of cells analyzed; error bar, s.e.m.; scale bar, 10μm. *P*-values were from *t*-test. ****, *P* ≤ 0.00005.

**Supplementary Figure 2.**
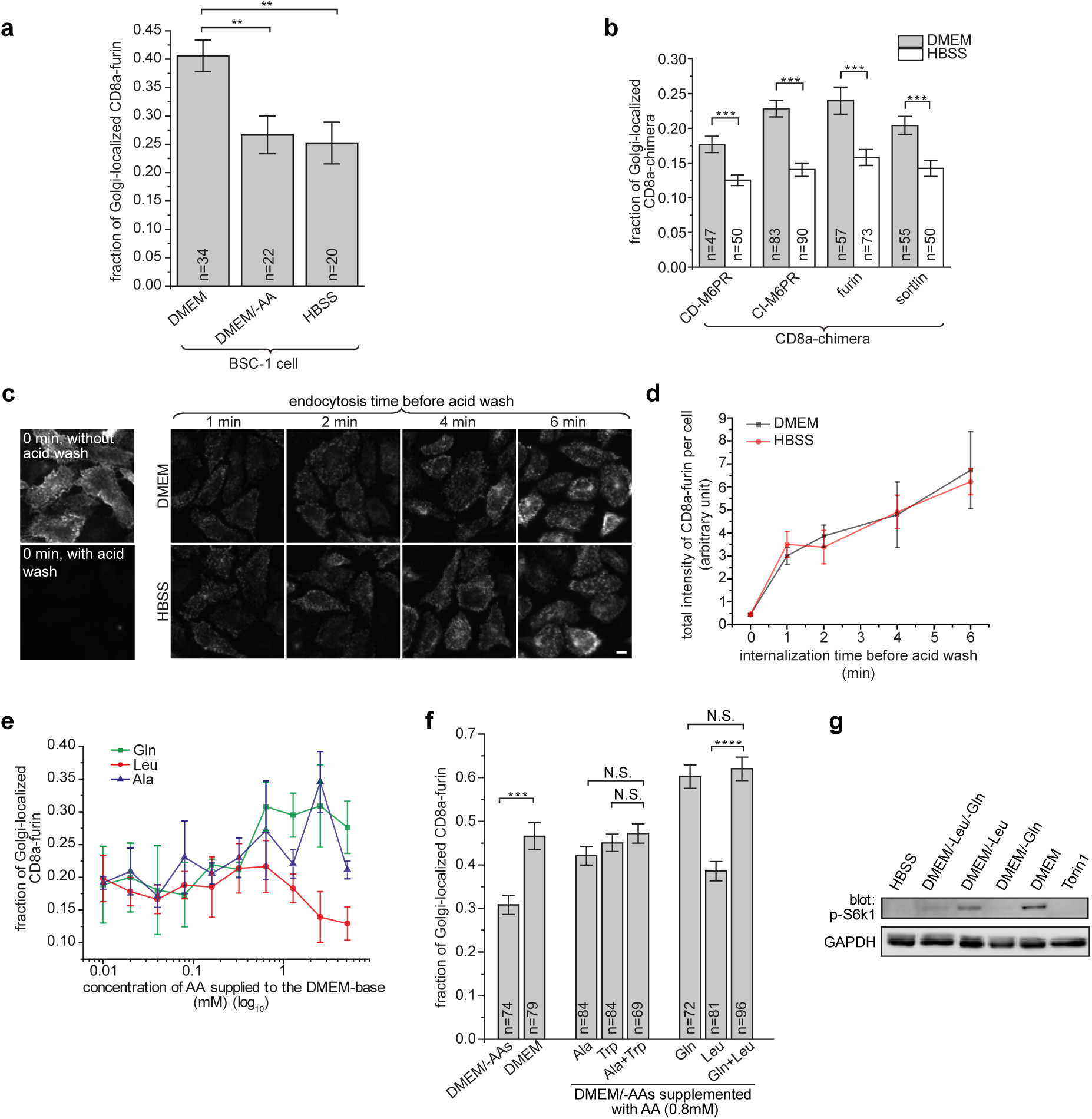
AAs stimulate endocytic trafficking to the Golgi. (**a**) BSC-1 cells transiently expressing CD8a-furin were incubated in HBSS for 2 h followed by surface-labeling using anti-CD8a antibody. Cells were subsequently incubated in indicated medium for 20 min before immuno-labeling for CD8a and endogenous giantin. The fraction of the Golgi-localized CD8a-furin is quantified. (**b**) Similar to **a** except that HeLa cells transiently expressing indicated CD8a-fused chimeras were used. (**c**,**d**) AAs do not significantly affect the endocytosis of CD8a-furin. HeLa cells expressing CD8a-furin were incubated in HBSS for 2 h followed by surface-labeling by anti-CD8a antibody. Next, labeled CD8a-furin was chased at 37 °C for indicated time before being subjected to acid wash on ice to remove the surface-bound antibody. Total cellular CD8a-furin was stained in **c** and the relative total CD8a intensity per cell was quantified in **d** as (total intensity of background-subtracted image)/(number of cells in the image). n=9 images were used for each time point. (**e**) HeLa cells stably expressing CD8a-furin were treated with DMEM/-AAs for 2 h followed by surface-labeling. The antibody-labeled CD8a-furin was subsequently chased for 20 min in DMEM/-AAs supplemented by indicated AA from 0.01 to 5.12 mM. After staining CD8a and giantin, cells were imaged and the Golgi-localized (*I_golgi_*) and total CD8a-furin intensity (*I_total_*) within each field of view were acquired. The fraction of Golgi-localized CD8a-furin is calculated as *I_golgi_*/*I_total_*. The plot incorporates n=4 (Ala and Gln) or 5 (Leu) randomly chosen fields of views with each field having ≥ 60 cells. (**f**) The effect of combining two AAs on the endocytic trafficking to the Golgi. Cells were treated as in **e** except with 0.8 mM indicated AAs. (**g**) Gln is essential for Leu to activate mTORC1 activity, as previously reported. HeLa cells were nutrient starved in HBSS for 2 h before treatment with indicated medium or Torin1 (in the complete medium) for 20 min. Endogenous phospho-S6K1 (p-S6K1) and GAPDH were blotted. Error bar in **a**, **b** and **f**, s.e.m.; error bar in **d** and **e**, s.d.; n, the number of cells analyzed; scale bar, 10 μm. *P*-values were from t-test. N.S., not significant (*P* > 0.05); **, *P* ≤ 0.005; ***, *P* ≤ 0.0005; ****, *P* ≤ 0.00005.

**Supplementary Figure 3.**
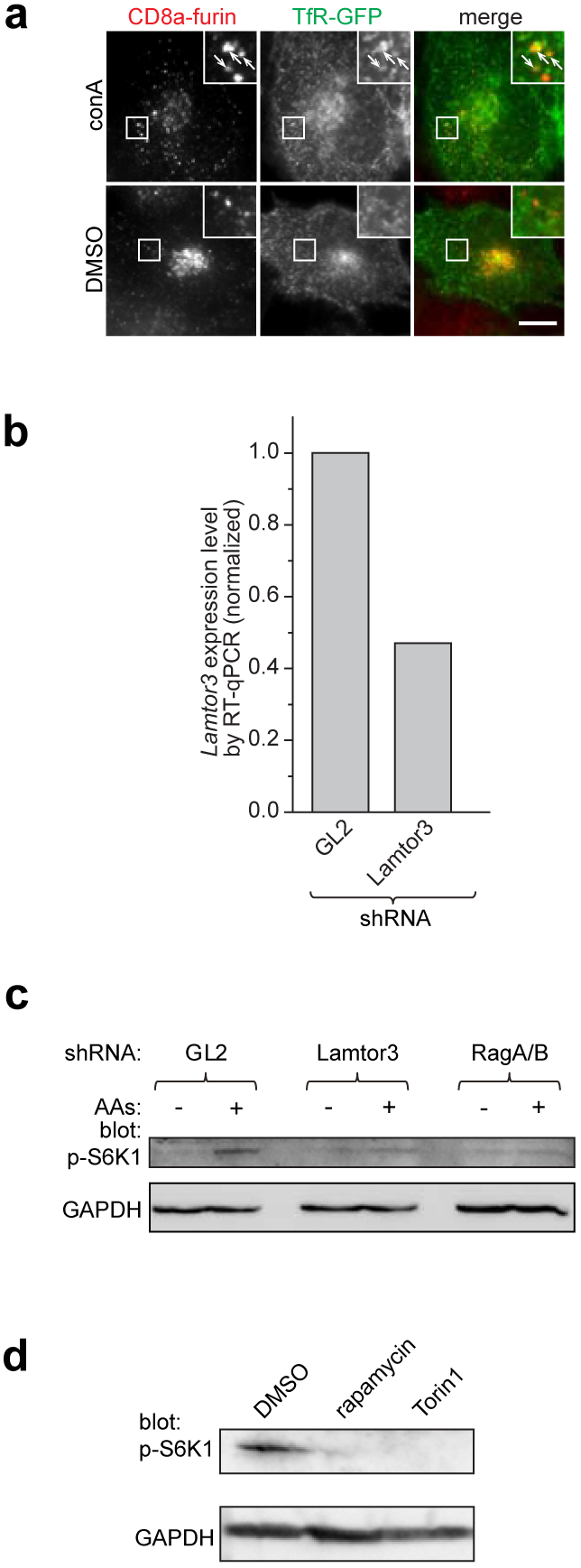
This figure corresponds to Fig. 3. (**a**) ConA induced the endosomal translocation of CD8a-furin. HeLa cells co-expressing CD8a-furin and TfR-GFP were incubated in HBSS for 2 h followed by DMEM for 20 min. 1% DMSO or 2.5 μM conA was present throughout both HBSS and DMEM incubation. Boxed regions are enlarged at the upper right corner to demonstrate the colocalization (indicated by arrows). (**b**) Confirmation of the knockdown of Lamtor3. HeLa cells were subjected to lentivirus-mediated knockdown using control shRNA (GL2) or shRNA targeting Lamtor3. The expression level of *Lamtor3* gene was subsequently quantified by RT-qPCR. (**c**) Depletion of Lamtor3 or RagA/B attenuated the AA-stimulated mTORC1 activity. The experiment was conducted as in Fig. 3c. (**d**) Rapamycin and Torin1 inhibited the AA-stimulated mTORC1 activity. The experiment was conducted similarly to Fig. 3g. Cell lysates were subjected to immuno-blotting for p-S6K1 and GAPDH.

**Supplementary Figure 4.**
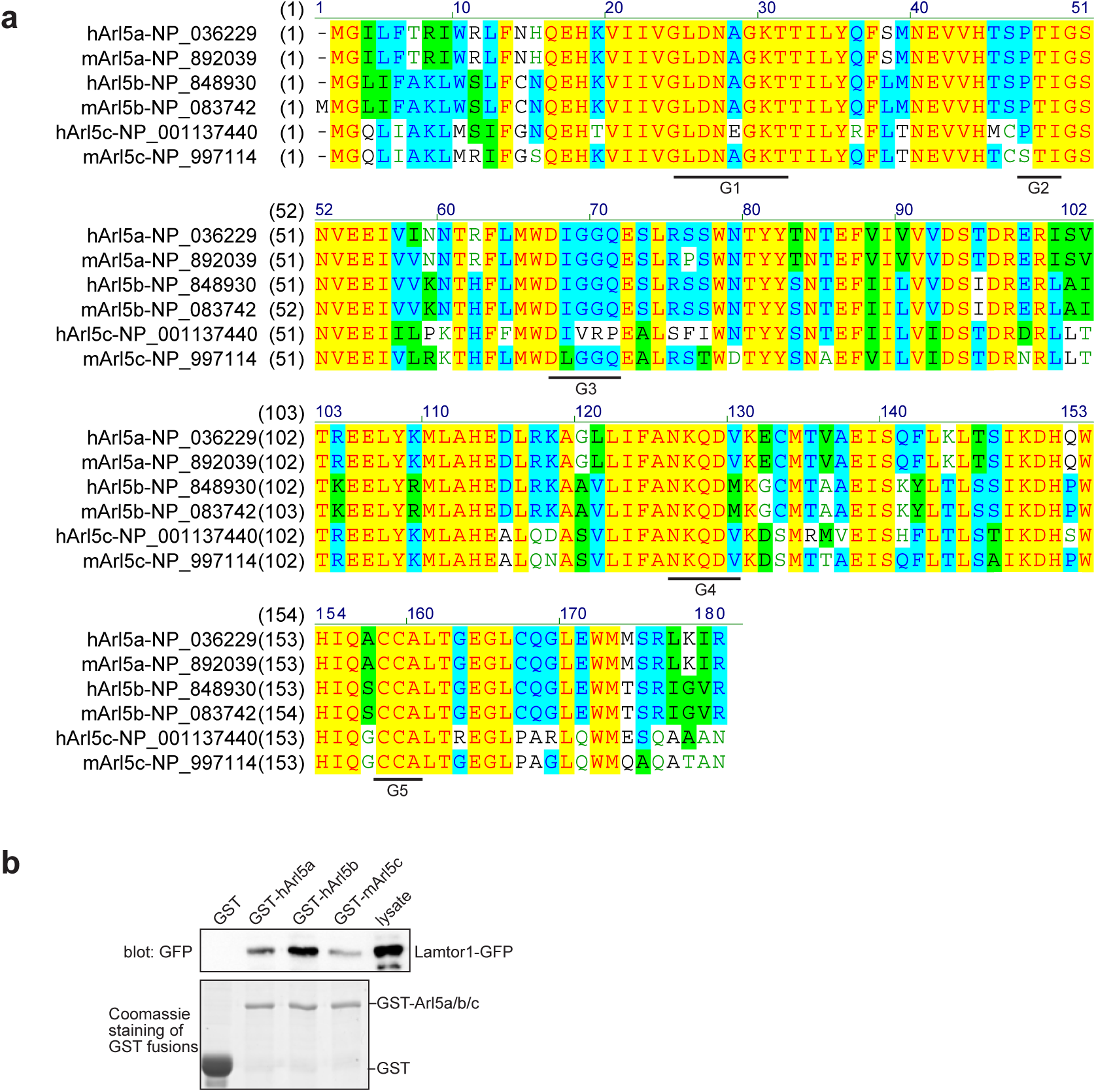
Arl5a and b are highly similar in primary sequence and both interact with Lamtor1. (**a**) Multiple sequence alignment of human and mouse Arl5a, b and c. The GenBank Accession number of each protein sequence is indicated. The five highly conserved guanine nucleotide binding motifs, G1-5 boxes, are underlined. The protein sequence of hArl5c is significantly different from that of Arl5a, Arl5b or mArl5c, especially at G3. The multiple alignment was conducted in Vector NTI (Invitrogen). (**b**) Human Arl5a and b and mouse Arl5c interact with Lamtor1. Bead-immobilized GST-fusions were incubated with cell lysate expressing Lamtor1-GFP and pull-downs were analyzed by immuno-blotting GFP-fusions. The prefix h and m denote human and mouse, respectively.

**Supplementary Figure 5.**
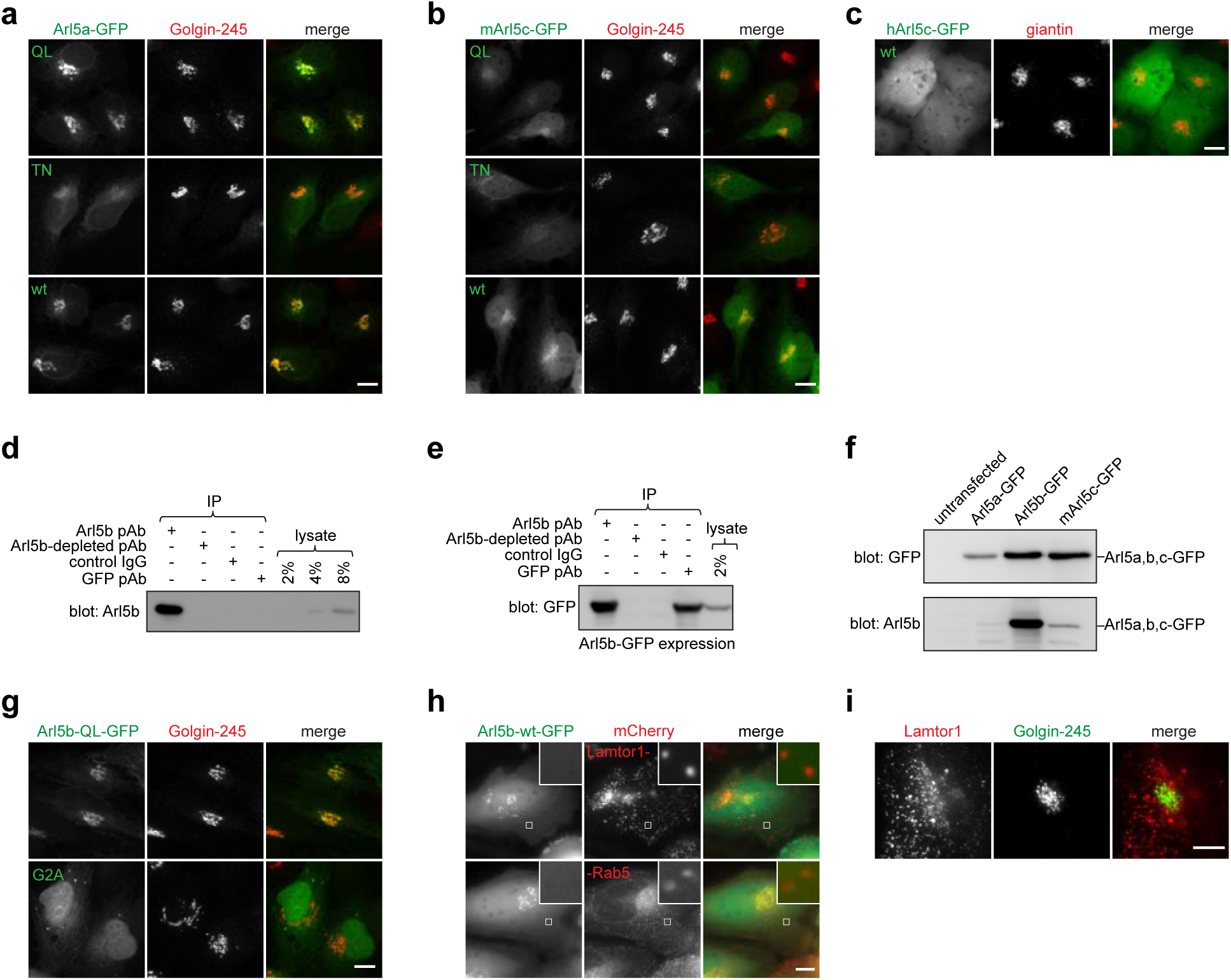
This figure corresponds to Fig. 5. (**a,b,c**) Arl5a and mArl5c, but not hArl5c, localize to the Golgi. HeLa cells transiently expressing indicated GFP-fusions were fixed and endogenous Golgin-245 or giantin was stained. The prefix h and m denote human and mouse, respectively. By default, Arl5 paralogs are from human. (**d,e,f**) Characterization of anti-Arl5b rabbit pAb. In **d**, HeLa cell lysates were subjected to co-IP by indicated antibodies and co-IPs were immuno-blotted by anti-Arl5b rabbit pAb. In **e**, HeLa cell lysates transiently expressing Arl5b-GFP were subjected to co-IP by indicated antibodies and co-IPs were analyzed by immuno-blotting GFP-fusions. To prepare Arl5b-depleted pAb, Arl5b pAb was incubated with sufficient amount of bead-immobilized GST-Arl5b and the supernatant was saved. In **f**, HeLa cell lysate transiently expressing GFP-tagged Arl5a, Arl5b or mArl5c was subjected to immuno-blotting by anti-GFP mAb and anti-Arl5b pAb, demonstrating that our anti-Arl5b pAb preferentially recognizes Arl5b. (**g**) N-terminal myristoylation is probably required for the Golgi localization of Arl5b. HeLa cells transiently expressing Arl5b-QL-GFP or Arl5b-QL-GFP harboring G2A mutation were fixed and endogenous Golgin-245 was stained. In Arl5b, Gly at position 2 is a potential myristoylation site. (**h**) Arl5b-wt does not significantly localize to the endosome or lysosome. Live imaging of HeLa cells expressing indicated proteins. (**i**) Lamtor1 does not localize to the Golgi. HeLa cells were stained for endogenous Lamtor1 and Golgin-245. Scale bar, 10 μm.

**Supplementary Figure 6.**
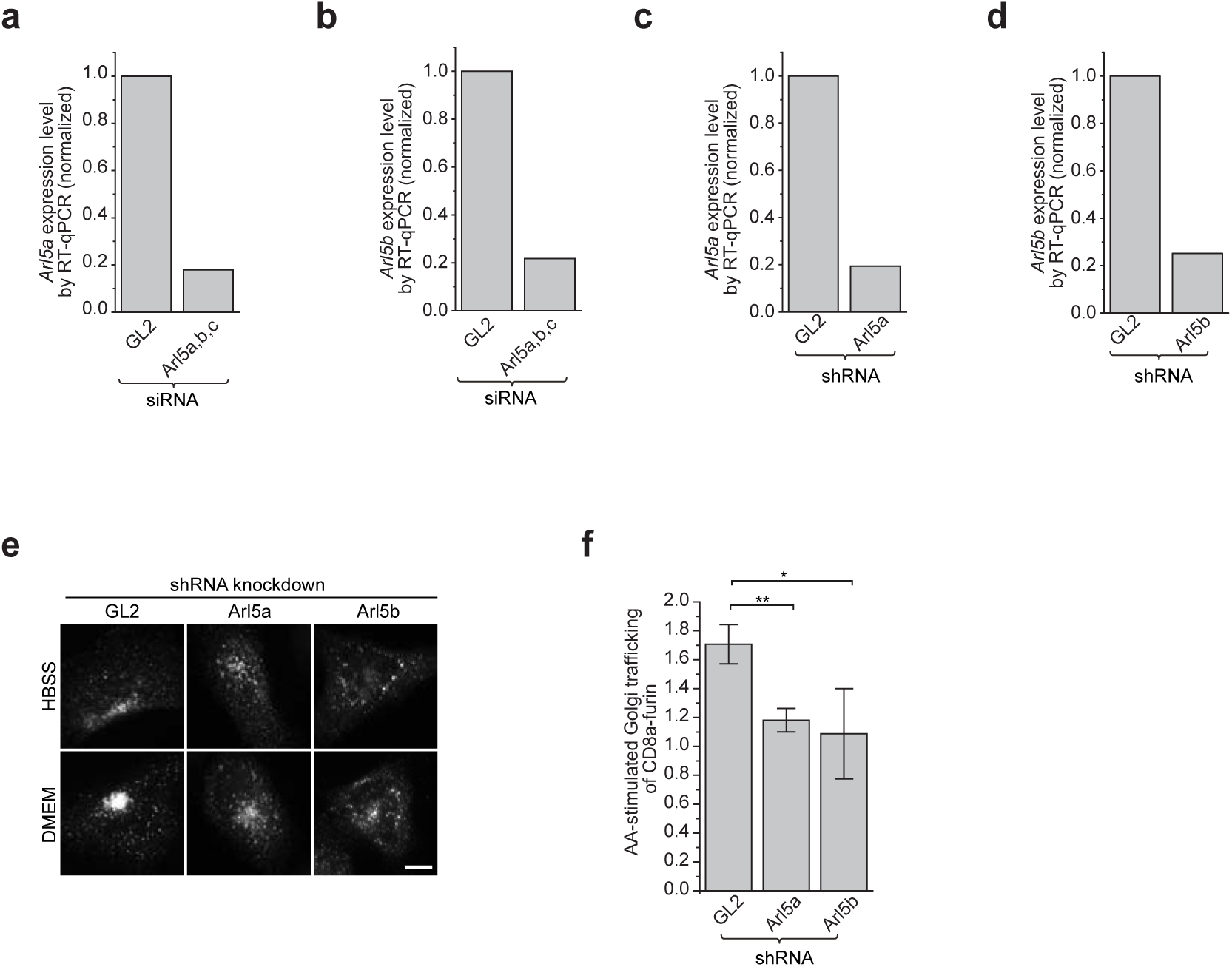
Arl5a and Arl5b are essential for the AA-stimulated endosome-to-Golgi trafficking of CD8a-furin. (**a,b**) HeLa cells were transfected with non-targeting control siRNA (GL2) or a mixture of siRNAs targeting Arl5a, b and c. The transcript expression level of Arl5a or b was quantified by RT-qPCR. The expression level was normalized by that of the corresponding control siRNA. (**c,d**) The control knockdown (GL2) or the knockdown of endogenous Arl5a or Arl5b by lentivirus-mediated transduction of corresponding shRNA. The transcript level was quantified as in **a** and **b**. In (**e,f**), HeLa cells were knocked down by lentivirus-mediated transduction of control shRNA (GL2) or shRNA targeting Arl5a or Arl5b. Cells were subsequently transfected to express CD8a-furin and treated with HBSS for 2 h followed by HBSS or DMEM for 20 min before immunofluorescence labeling of CD8a-furin and giantin (not shown). The AA-stimulated Golgi trafficking of CD8a-furin was calculated for each shRNA. The displayed value is the mean of n=3 independent experiments with each experiment analyzing ≥ 17 cells. Error bar, s.d.; scale bar, 10 μm. *P*-values were from *t*-test. *, *P* ≤ 0.05; **, *P* ≤ 0.005.

